# Induction of PARP7 Creates a Vulnerability for Growth Inhibition by RBN2397 in Prostate Cancer Cells

**DOI:** 10.1101/2022.09.02.506406

**Authors:** Chunsong Yang, Krzysztof Wierbiłowicz, Natalia M Dworak, Song Yi Bae, Sachi B. Tengse, Nicki Abianeh, Justin M. Drake, Tarek Abbas, Aakrosh Ratan, David Wotton, Bryce M Paschal

## Abstract

The ADP-ribosyltransferase PARP7 modulates protein function by conjugating ADP-ribose to the side chains of acceptor amino acids. PARP7 has been shown to affect gene expression in prostate cancer cells and certain other cell types by mechanisms that include transcription factor ADP-ribosylation. Here, we use a recently developed catalytic inhibitor to PARP7, RBN2397, to study the effects of PARP7 inhibition in androgen receptor-positive and androgen receptor-negative prostate cancer cells. We find that RBN2397 has nanomolar potency for inhibiting androgen-induced ADP-ribosylation of the androgen receptor. RBN2397 inhibits the growth of prostate cancer cells in culture when cells are treated with ligands that activate the androgen receptor, or the aryl hydrocarbon receptor, and induce PARP7 expression. We show that the growth inhibitory effects of RBN2397 are distinct from its enhancement of interferon signaling recently shown to promote tumor immunogenicity. RBN2397 treatment also induces trapping of PARP7 in a detergent-resistant fraction within the nucleus, which is reminiscent of how inhibitors such as Talazoparib affect PARP1 fractionation. Because PARP7 is expressed in AR negative metastatic tumors and RBN2397 can affect cancer cells through multiple mechanisms, PARP7 may be an actionable target in advanced prostate cancer.

**Significance:** RBN2397 is a potent and selective inhibitor of PARP7 that reduces the growth of prostate cancer cells, including a model for treatment-emergent neuroendocrine prostate cancer. RBN2397 induces PARP7 trapping on chromatin, suggesting its mechanism of action might be similar to clinically-used PARP1 inhibitors.

## INTRODUCTION

The PARP family of enzymes contributes to a variety of cellular pathways, many of which occur in the nucleus and involve events associated with the regulation of chromatin structure, transcription, and DNA damage signaling and DNA repair[1, 2]. PARP enzymes contain a conserved catalytic domain that uses NAD^+^ as a cofactor for ADP-ribose conjugation to the side chains of acceptor amino acids, and in the case of enzymes that generate polymers, conjugation to ADP-ribose itself[3]. The founding member of the PARP family, poly-ADP-ribosyltransferase 1 (PARP1) has been a major focus in the field since its discovery. Detailed, mechanistic analysis of PARP1 has been complemented by the development of highly specific compounds that compete for NAD^+^ binding, and as a consequence, inhibit PARP1 enzymatic function[4]. Clinical trials with PARP1 inhibitors such as Olaparib have shown patient benefit in ovarian, breast, and prostate cancer, particularly in tumors that harbor mutations in DNA repair genes, notably *BRCA1/2* and *ATM*[5].

Most PARP family members mediate a single round of ADP-ribose attachment and are categorized functionally as mono-ADP-ribosyltransferases[3]. Mono-ADP-ribosylation provides a post-translational mechanism for modulating protein function, likely in a reversible manner since cells encode hydrolases that can remove ADP-ribose from amino acids[6]. TIPARP/PARP7, a mono-ADP-ribosyltransferase expressed in multiple cell and tissue types, was characterized as a key effector of signaling and gene expression mediated by the aryl hydrocarbon receptor (AHR) in the context of detoxification pathways in liver[7]. Our group identified PARP7 in a prostate cancer cell signaling pathway that controls assembly of a multi-subunit complex containing the androgen receptor (AR) [8].

In this nuclear pathway, PARP7 “writes” ADP-ribose onto cysteine (Cys) residues in AR. The ADP-ribosyl-Cys sites are subsequently “read” by macrodomains in PARP9; this provides a highly selective mechanism for assembling a complex that contains AR, PARP9, and the ubiquitin E3 ligase DTX3L. Depletion of DTX3L affects the expression of a subset of AR-regulated genes[8], suggesting PARP7 and assembly of the complex has a regulatory role for AR-mediated transcription. PARP7 has been functionally linked to other transcription factors including the estrogen receptor and liver X receptors [9, 10]. It seems plausible that PARP7 contributes to a variety of gene expression pathways through mechanisms that include, but may not be limited to, transcription factor ADP-ribosylation.

PARP7 also exerts an effect on transcription through a signaling-based mechanism involving the kinase, TBK1. PARP7 negatively regulates TBK1 kinase activity, which restrains phosphorylation and activation of the transcription factor IRF3[11]. In this pathway, PARP7 was proposed to serve as a brake for sensing cytosolic nucleic acids that trigger Type I IFn signaling [12]. Ribon Therapeutics developed RBN2397 as a first-in-class mono-ADP-ribosyltransferase inhibitor, and showed that it blocks PARP7 negative regulation of TBK1[12]. Treating lung cancer models with RBN2397 promotes TBK1 phosphorylation of IRF3, restores Type I IFN signaling, and enhances tumor immunogenicity [12]. These data provide the rationale for targeting PARP7 with RBN2397 and its evaluation in clinical trials (NCT04053673).

Here, we explore RBN2397 as a PARP7 inhibitor in prostate cancer cells. Consistent with a previous report[12], RBN2397 has selectivity for PARP7 versus other PARP family members in ADP-ribosylation assays using core histones as substrates. In cells, we show that RBN2397 has nanomolar potency for inhibiting PARP7 ADP-ribosylation of AR. We determined that RBN2397 inhibits the growth of prostate cancer cells dependent on transcriptional induction of PARP7, achievable by treating cells with androgen to activate AR, or with ligands that activate AHR. RBN2397 exerts a nuclear trapping effect on PARP7, similar in nature to the effects of clinically-used inhibitors to PARP1[13, 14]. Finally, we used chemical inhibitors to TBK1 and JAK1/2 to show that RBN2397 inhibition of prostate cancer cell growth can be distinguished from the RBN2397 effect on PARP7 regulation of Type I IFN signaling. The available data indicate that PARP7 inhibition with RBN2397 exerts effects on cancer cells through multiple mechanisms.

## MATERIALS AND METHODS

### Drugs and chemicals

Synthetic androgen methyltrienolone (R1881, Perkin-Elmer NLP005005MG), PARP7 inhibitor RBN2397 (RBN, Medchemexpress LLC HY-136174 or DCCHEMICALS DC31069), PARP1/2 inhibitor Veliparib (Selleck Chemical LLC S100410MG), 10-CL-BBQ (BBQ, Tocris Bioscience 63-215), FICZ (Medchemexpress LLC HY-12451), JAK1/2 inhibitor Ruxolitinib (Ruxo, Selleck Chemical LLC S13785MG), TBK1 inhibitor GSK8612 (GSK, Medchemexpress LLC HY-111941), IFNαA (MilliporeSigma IF007), proteasome inhibitor MG132 (Caymen Chemical 1001262810), Cycloheximide (MP Biomedicals 0219452701), (3-(4,5-dimethylthiazol-2-yl)- 2,5-diphenyltetrazolium bromide (MTT, Tocris Bioscience 5224), 50x B27 supplement (Thermo Fisher Scientific 17504044), recombinant human βFGF (Peprotech 100-18b) and recombinant human EGF (Peprotech AF-100-15).

### Cell culture

Cell lines were obtained from ATCC: PC3 (CRL-1435), VCaP (CRL-2876), CWR22Rv1 (CRL-2505), DU145 (CRL-HTB-81), C4-2b (CRL-3315), HEK293T (CRL-3216) and NCI-H660 (CRL-5813). Cell lines were mycoplasma-free upon receipt from ATCC. Cell lines were typically grown up to 20-30 passages, and are male-derived. The following growth media used: RPMI1640 + 5% FBS (PC3, DU145, and derivatives), DMEM + 10% FBS (VCaP), RPMI1640 + 10% FBS (CWR22Rv1), DMEM + 5% FBS (HEK293T) and Advanced DMEM/F12 (Gibco) + 1× B27 Supplement (Gibco) + 10 ng/mL EGF (PeproTech) + 10 ng/mL bFGF (PeproTech) + 1× Glutamax (Life Technologies) (NCI-H660). All media were supplemented with 1% Pen/Strep.

### Growth assays

Cells (100 μl/well, typically seeded at 1:40 in growth media combined with drugs R1881, RBN2397, BBQ, FICZ, Ruxo, GSK and IFNαA were grown at 37°C in 96-well plates in eight biological replicates, with media changes every 2 days, until control cells reached sufficient density. The cells were washed once with PBS, incubated at 37°C for 2 hours with 100 μl of phenol-red free DMEM (for VCaP) or RPMI (for others) + 2% FBS + 0.25 mg/ml MTT, followed with medium removal, extraction with 100 μl of DMSO, and measurements on Synergy HT plate reader (BioTek). NCI-H660 cells (5,000 cells/well, n = 5) were treated with RBN2397, BBQ and FICZ as indicated at 37°C in 96-well plates for 5 days prior to measuring viability using WST reagent (Takara) following the manufacturer’s protocol. The plates were read on Infinite M1000 Pro microplate reader (Tecan). The data was analyzed with Prism (GraphPad). Error bars represent standard deviation. P value **** <0.0001; *** <0.001; ** <0.01; * <0.05; ns, not significant.

For live cell microscopy, cells were drug-treated for 6 days (slower growing VCaP and CWR22Rv1) or 3 days (faster growing PC3-AR and PC3). Images were taken under the EVOS Cell Imaging System (Thermo Fisher). NCI-H660 cells were drug-treated for 5 days and images were taken under a microscope using ZEN microscopy software (Carl Zeiss Microscopy). For dead cell measurements, cells were treated with trypsin, collected by centrifugation (2,000x RPM, 3 minutes). Cells were re-suspended in equal volume of PBS and Trypan Blue Stain 0.4% (Invitrogen T10282), and counted on Countess II FL (Life Technologies).

### Cell cycle and apoptosis assays

Cells were drug-treated for 48 hrs (slower growing VCaP and CWR22Rv1) or 24 hrs (faster growing PC3-AR and PC3). Cells were trypsinized, collected by centrifugation, washed once with cold PBS, resuspended in 0.5 ml of PBS, mixed to 4.5 ml of cold 70% ethanol, and stored on ice until usage. Prior to staining, cells were collected by centrifugation and washed with PBS. Cells (~3.5 x 10^5^) were resuspended in 0.1 ml of staining solution (PBS + 0.1% Triton X-100 (TX-100) + 80 μg/ml of Propidium Iodide + 0.2 mg/ml of RNase A), incubated at 37°C for 40 minutes, diluted with 0.4 ml of PBS + 0.1% TX-100, measured using an Attune NxT Cytometer (Thermo Fisher), and quantified with FCS Express 7 (De Novo Software). For the apoptosis assay, NCI-H660 cells were drug-treated for 5 days, and dispersed into single cells by trypsin treatment and pipetting. Cells were then filtered through 35 μm nylon mesh and stained with Annexin V-FITC and 7-AAD (Biolegend). 50,000 cells per sample were collected for analysis using a LSRII flow cytometer (BD Biosciences) and the distribution of cells were quantified with FlowJo (TreeStar).

### siRNA

PC3-AR cells were transfected with 20 nM of Negative Control siRNA #1 (*siCon*, Ambion AM4635), *siPARP7* (sense strand 5’-AAUACUCUCAUCGAACGGAAGTT-3’) or *si-p21* (sense strand 5-AACAUACUGGCCUGGACUG-3’) using Lipofectamine RNAiMAX (Invitrogen 56532). After 24 hrs of transfection, cells were re-plated into 96-well plates for cell growth assays. Percentages of cell growth inhibition at various RBN2397 concentration were compared between *siPARP7* (or *si-p21*) and *siCon*, with p value **** <0.0001; *** <0.001; ** <0.01; * <0.05. To confirm *PARP7* (or *p21*) knockdown, cells (24 hrs post-transfection) were re-plated into 6-well plates and after 48 hrs, cells were treated with R1881 and RBN2397 for 24 hrs. Western blotting was used to measure relative protein expression.

### Lentiviruses and cell line generation

PC3-AR, PC3-AR(*HA-PARP7*) and PC3-AR(*shGFP* or *shPARP7*) have been previously described[8].

Lentiviral plasmid *shPARP7* (or *shGFP* control, or pLH3/*AviTag-PARP7*) and two accessory plasmids, pMD2g and psPAX2 (0.5x amount of the lentiviral plasmid), were transfected into HEK293T cells with transfection reagent ViaFect (Promega PRE4981). After 16 hrs of transfection, growth media was changed to DMEM + 35% FBS, and cells grew another 24 hrs. The growth media containing the lentiviruses was transferred to a conical tube. Cell debris was removed with centrifugation at 1,000 RPM. The virus solution was passed through 0.4 μm filter, concentrated with Lenti-X Concentrator (Takara 631231). CWR22Rv1 or C4-2b cells were infected with virus for 24 hrs in the presence of 8 μg/ml of polybrene in the growth medium. Cells were then changed into fresh growth media, grown for 2-3 doubling, and then selected with 2 μg/ml of puromycin.

### *CDKN1A* deletion

A pair of sgRNAs targeting nonoverlapping sequences within Exon 2 of the *CDKN1A* gene, encoding p21, were designed using Benchling, and cloned into the CAS9-expresssing pX330 expression vector (Addgene #42230). The following sgRNAs (sense strand) were used; sg-*p21*-1: 5’-GCGCCATGTCAGAACCGGCT-3’, ‘ sg-*p21*-2: 5’-TTAGCGCATCACAGTCGCGG-3’. PC3-AR cells were transfected with the two CAS9-expressing plasmids (sg-*p21*-1 and sg-*p21*-2; 2.0 μg each), along with 0.5 μg pMSCVpuro vector (Clontech). After selection, single clones were isolated, serially diluted, expanded, and genotyped. Genotyping was performed by PCR amplification of the targeted *p21* locus with primers flanking the two predicted CAS9 cleavage sites followed by Sanger sequencing. The following primers were used to amplify a 339 bp sequence spanning the two sgRNA target sites: p21-F: 5’-TCACCTGAGGTGACACAGCAAAGC-3’, p21-R: 5’-GGCCCCGTGGGAAGGTAGAGCTT-3’. Immunoblotting of the clonal cells with antip21 antibody was used to confirm loss of protein expression.

### Antibodies

Rabbit antibodies to AR (1 μg/ml) and PARP7 (1 μg/ml) have been described[8, 15]. Commercial antibodies were used against TUBULIN (1:10,000–20,000, mouse mAb TUB-1A2, Sigma T9028), HA tag (1:1,000, mouse mAb 16b12, Covance A488-101L, also used for IP), p21 (1 μg/ml, mouse mAb SX118, BD Pharmingen 550827), AHR (1 μg/ml, Rabbit mAb JM34-10, Thermo Fisher Scientific MA5-32576), CYCLIN B1 (1:10,000, rabbit mAb Y106, Epitomics 1495-1), CYCLIN A (1:500, mouse mAb 6E6, Leica Microsystems NCL-CYCLIN A), STAT1α (1:1,000, rabbit mAb EPR4407, abcam ab109320), p-STAT1(Tyr701)(1:1,000, rabbit, Cell Signaling 9171), LAMIN A/C (1:40, mouse mAb JOL2, Millipore MAB3211), Alexa Fluor 680 donkey anti-Rabbit IgG(H + L) (1:20,000, Invitrogen A10043), IRDye800-conjugated anti-mouse IgG(H&L) (1:20,000, Rockland 610-132-121), and AR (mouse mAb G122-434, BD Pharmingen 554225, used for IP). PARP7 and AR (or AHR) were detected sequentially by using the aforementioned antibodies and detected on Odyssey CLx (LI-COR).

### AR ADP-ribosylation detection

PC3-AR cells were grown in the growth media supplemented with 2 nM R1881 +/- RBN2397 for 16-20 hours. After media removal, cells were lysed in 1xSDS loading buffer (200mM Tris-HCl pH 6.8, 2% SDS, 10% glycerol and 3% β-mercaptoethanol), and the lysates were heated at 95°C for 5 min, followed with sonication, centrifugation, SDS-PAGE separation, and nitrocellulose membrane transfer. The membrane was blocked in PBST + 5% nonfat milk at room temperature for 45 minutes, followed with overnight 4°C incubation with 1 μg/ml Fl-Af521 [16], room temperature PBST washing (5 minutes each for 5 times), and detection on ODYSSEY CLx (LI-COR).

### Protein half-life measurement

PC3-AR(*HA-PARP7*) cells, with or without 1 hr of 10 nM RBN2397 pre-treatment, were incubated with 0.1 mg/ml of cycloheximide for indicated time periods. After media aspiration, cells were lysed in 1xSDS-PAGE loading buffer, followed with heating, sonication, SDS-PAGE, HA-PARP7 Western Blot (with TUBULIN as control), and quantification on Odyssey CLx (LI-COR).

### PARP activity assays

The assays were conducted by BPS Bioscience (San Diego, CA) in duplicates (PARP7) or triplicates (PARP family) at room temperature for 2 hrs with reaction mixture containing PARP, inhibitors, β-NAD^+^, and Biotin-β-NAD^+^ in 96-well plates pre-coated with histone substrates. After enzymatic reactions, 50 μl of Streptavidin-horseradish peroxidase (prepared with Blocking Buffer) was added to each well and the plate was incubated at room temperature for an additional 30 min. The wells were washed again and 100 μl ELISA ECL substrate was added to each well. Luminescence was measured using a BioTek SynergyTM 2 microplate reader.

### Cell fractionation

All procedures were performed at 4°C. Cells were PBS washed once, scraped in Buffer A (20 mM Tris-HCl pH 7.5, 100 mM NaCl, 1 mM PMSF and 1 μM Veliparib), and collected by centrifugation (950×*g*, 5 min). Cells were lysed in Buffer B (20 mM Tris-HCl pH 7.5, 100 mM NaCl, 0.5% TX-100, 1 mM PMSF, 2 mM DTT, 2 mM EDTA, 1 μM Veliparib, and 5 μg/ml each of aprotinin/leupeptin/pepstatin) for 20 min. Supernatants were collected after centrifugation (16,800×*g*, 20 min) and deemed as the TX-100 soluble fraction. The remaining pellets were washed once with Buffer B, resuspended in 1xSDS loading Buffer C (200mM Tris-HCl pH 6.8, 10% glycerol, 2% SDS and 3% β-mercaptoethanol), sonicated, subjected to centrifugation (16,800xg, 20 min) to obtain the supernatant which was deemed the TX-100 insoluble fraction.

For immunoprecipitation, the TX-100 soluble fraction was subjected to 4 μg antibody/2 μl packed Protein G magnetic beads per IP for binding for 3-4 hrs. The beads were separated under magnetic fields, followed with 5 washes with Buffer D (20 mM Tris-HCl pH 7.5, 100 mM NaCl, 0.1% TX-100, 1 mM DTT, 0.1 mM EDTA, 1 μM Veliparib, and 1 μg/ml each of aprotinin/leupeptin/pepstatin), and then resuspended in 1xSDS loading Buffer C.

### AHR Transcription Assay

Cells were grown in triplicates in the presence or absence of AHR ligands BBQ (100 nM) and FICZ (1 μM) for 18 hrs. RNA isolation was done using the Qiashredder (Qiagen 79656) and RNeasy Mini Kit (Qiagen 74106), followed with iScript cDNA synthesis (Bio-Rad 1708891) and real-time qPCR using primer sets (CYP1A1: CAACCCTTCCCTGAATGCCT and GCTTCTCCTGACAGTGCTCA; CYP1B1: AACGTACCGGCCACTATCAC and TCACCCATACAAGGCAGACG) and 2x SensiMix SYBR & Fluorescein Mastermix (Bioline QT615-05) on the iCycler (Bio-Rad).

### Immunofluorescence (IF) and Confocal Microscopy

Cells were washed with PBS, fixed at room temperature (RT) for 20 min using the 10% neutral buffered formalin (Thermo Scientific 5701), washed with PBS, and then permeabilized at RT for 20 min with PBS + 0.2% TX-100; alternatively, cells washed with cold PBS, permeabilized at 4°C for 20 min using extraction buffer (20mM Tris-HCl pH 7.5, 100mM NaCl, 2mM EDTA, 2mM DTT, 1mM PMSF, 5μg/ml each of aprotinin/leupeptin/pepstatin), washed with the cold extraction buffer followed with cold PBS, and fixed at RT for 20 min using the 10% neutral buffered formalin. After above treatments, cells were washed 3x with PBST, followed with a routine blocking (PBST + 5% BSA, RT, 60 min), primary antibody incubation (in PBST + 1% BSA, at RT for 2 hrs), 3x PBST washing, fluorescently labeled secondary antibody incubation (in PBST + 1% BSA, at RT for 2 hrs), DAPI stained (2μg/ml in PBS, at RT for over 14 seconds), milli Q water washed, mounted on slides using Fluoro-Gel (Electron Microscopy Sciences 17985-10), and sealed with a nail polish (Electron Microscopy Sciences 72180). Images shown in the figures were acquired by confocal microscopy (LSM 880; Carl Zeiss) equipped with a 63x, 1.4NA oil immersion objective and ZEN software (Carl Zeiss). All IF images shown were processed in parallel with Photoshop (Adobe), ImageJ and figures were assembled using Illustrator (Adobe).

#### RNA-seq data analysis

All of the data used in this study except GSVA and PARP7 gene amplification analysis was obtained from recount3. Primary tumor data for prostate (PRAD) and lung (LUAD) used for GSVA analysis was generated by the TCGA Research Network: https://www.cancer.gov/tcga.” TCGA PRAD RNA-seq dataset was prefiltered to keep only primary tumor and ffpe samples were removed (n=497). The metastatic castration-resistant prostate cancer (mCRPC) dataset (accn: SRP183532) from Fred Hutchinson Cancer Research Center [17] was prefiltered to keep only patient samples (n=98). Samples were divided into *AR*+ and *AR*-groups based on phenotypes evaluated in the original study[17]. For normal prostate analysis, we used the GTEx PROSTATE dataset (n=263). The Genotype-Tissue Expression (GTEx) Project was supported by the Common Fund of the Office of the Director of the National Institutes of Health, and by NCI, NHGRI, NHLBI, NIDA, NIMH, and NINDS. To assess the *PARP7* gene expression level cutoff we used our previously generated RNA-seq dataset (accn: SRP162931)[18], which also was obtained from recount3. An additional primary prostate cancer tumor data set (accn: SRP163173)[19] is from Netherlands Cancer Institute. For *PARP7* expression level comparison, we used counts per million (CPM) values calculated with the edgeR package in R. For correlation analysis, we used Spearman correlation on log2 transformed Transcripts Per Million (TPM) and Fragments Per Kilobase Million (FPKM) normalized data. For GSVA analysis [20], we used primary tumor samples from LUAD and PRAD TCGA cohorts. Samples were divided into high *PARP7* expressing (top quartile) and low *PARP7* expressing (bottom quartile). GSVA analysis was done using the GSVA Bioconductor R package for the MSigDB HALLMARK_INTERFERON_ALPHA_RES PONSE and HALLMARK_ANDROGEN_RESPONSE gene sets. We acknowledge our use of Molecular Signature Database (MSigDB)[21]. The data used for PARP7 gene amplification analysis was acquired from cBioPortal (https://www.cbioportal.org/). The metastatic prostate cancer tumor data set used in this analysis is from Stand Up to Cancer/Prostate Cancer Foundation (SU2C-PCF) International Prostate Cancer Dream Team consortium[22].

### Statistical analysis

Cell growth assays involved 8 biological replicates for each condition and analysis using two-tailed paired t-test. In RNA-seq analysis, Wilcoxon rank-sum test was used for comparisons between two groups and Kruskal-Wallis test for comparisons between more than two groups. Spearman correlations were calculated to assess all correlations. A p-value of below 0.05 was considered statistically significant.

### Data availability

The Genotype-Tissue Expression (GTEx) PROSTATE RNA-seq dataset, The Cancer Genome Atlas Prostate Adenocarcinoma (TCGA-PRAD) RNA-seq dataset, the metastatic castration-resistant prostate cancer (mCRPC) RNA-seq dataset from Fred Hutchinson Cancer Research Center (accn: SRP183532)[17], the primary prostate cancer tumor RNA-seq dataset from Netherlands Cancer Institute (accn: SRP163173)[19] and our previously generated VCaP cell RNA-seq dataset (accn: SRP162931)[18] were obtained from recount3 project (https://rna.recount.bio/) which uniformly reprocesses publicly available RNA-seq datasets from GTEx, TCGA and Sequence Read Archive (SRA) using a Monorail analysis pipeline.

The metastatic prostate cancer RNA-seq and Copy Number Alteration (CNA) datasets from Stand Up to Cancer/Prostate Cancer Foundation (SU2C-PCF) International Prostate Cancer Dream Team consortium and the TCGA-PRAD Copy Number Alteration (CNA) dataset come from cBioPortal (https://www.cbioportal.org/).

## RESULTS

RBN2397 reduces the growth of lung cancer cells by inhibiting PARP7, increasing IFN signaling, and improving tumor immunogenicity[12]. We evaluated whether RBN2397 has potential utility in prostate cancer using biochemical approaches and cell growth assays in AR-positive (AR+) and AR-negative (AR-) prostate cancer lines. In previous work we reported that PARP7 directly ADP-ribosylates eleven Cys residues in AR[8]. AR ADP-ribosylation is induced by androgen in prostate cancer cells because the *PARP7* gene is a direct target of AR, and because PARP7 selectively ADP-ribosylates the agonist conformation of AR [8, 23]. Androgen-induced ADP-ribosylation of AR in cells and detection with fluorescently-labeled Af1521 (Fl-Af521) on blots [16] provides a sensitive assay to assess the effect of chemical inhibition of PARP7 in cells. To this end, we treated PC3-AR cells with synthetic androgen (R1881) to induce *PARP7* transcription and AR-ADP-ribosylation, in the absence and presence of the PARP7 inhibitor RBN2397. We found that AR ADP-ribosylation was essentially eliminated by 3 nM RBN2397 (**Fig. 1A**). RBN2397 treatment prevents assembly of AR-DTX3L/PARP9 complexes (**Fig. 1B**), which was anticipated since PARP7 ADP-ribosylates Cys residues read by PARP9[8]. RBN2397 also inhibited androgen-induced AR ADP-ribosylation in VCaP cells (**Fig. 1C**). The selectivity of RBN2397 for PARP7 versus the PARP family was examined previously with biophysical assays that measure competitive binding with NAD^+^, which is the basis of how RBN2397 inhibits PARP7 enzyme function[12]. We measured the effect of RBN2397 on substrate ADP-ribosylation using recombinant PARP7 and PARP1 and a mixture of Histone H2A and H2B proteins to determine EC_50_ values. In vitro, the EC_50_ of RBN2397 for PARP7 is approximately 7.6 nM, whereas the EC_50_ of RBN2397 for PARP1 is 110 nM (**Fig. 1D**). In the presence of 7.6 nM RBN2397, there is relatively little inhibition of other PARP enzymes, including family members with RNA levels in prostate tumors that well exceed that of PARP7 (**Fig. 1E**).

**Fig.1.**
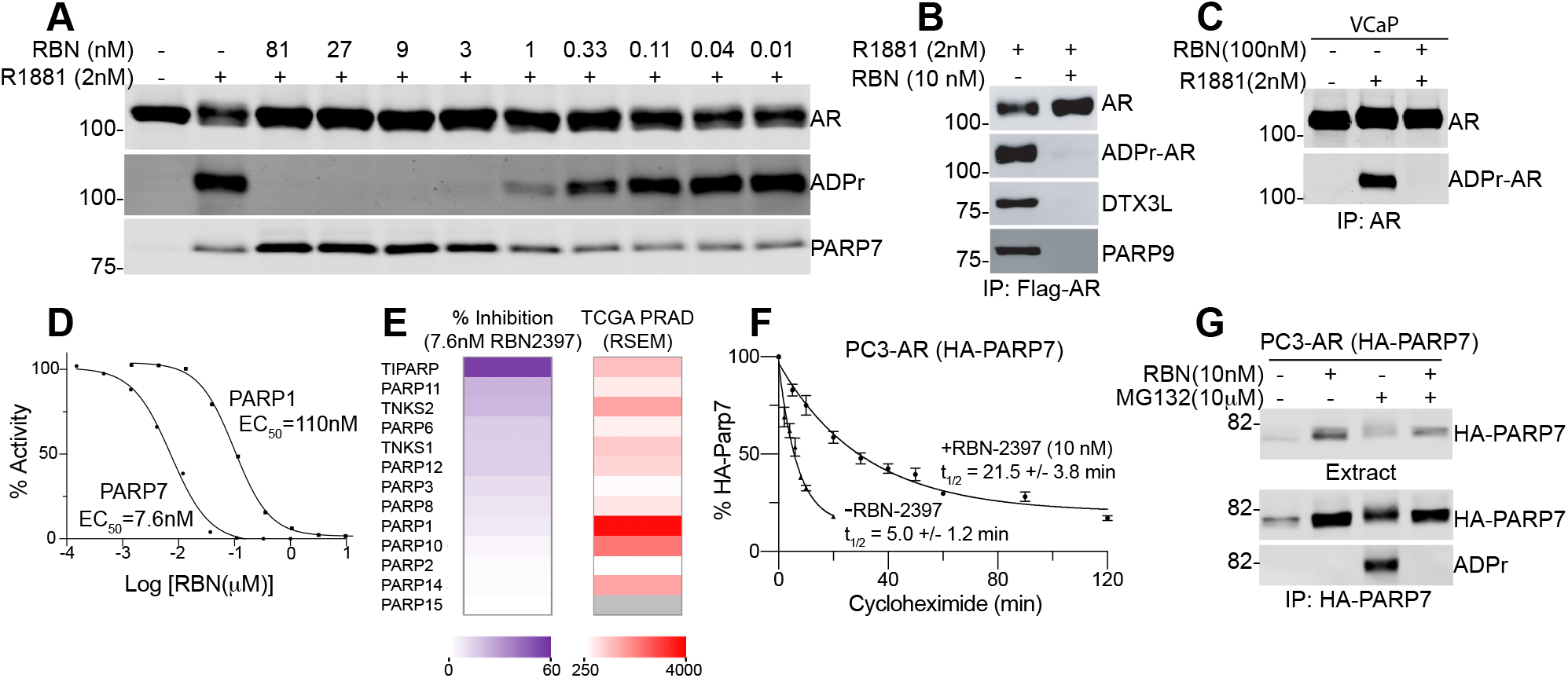
NAD^+^ competitive inhibitor RBN2397 blocks PARP7-mediated ADP-ribosylation of the androgen receptor (AR) in cells. (A) Effect of RBN2397 (12 pM to 81 nM) on androgen-induced AR ADP-ribosylation in PC3-AR cells treated for 17 hrs. AR and PARP7 were detected using Western blotting method. ADPr-AR was detected using Fl-Af521. (B) RBN2397 inhibits androgen-dependent AR ADP-ribosylation and AR/DTX3L/PARP9 complex formation in PC3-AR cells co-treated with R1881 for 17 hrs. Flag-AR was immunoprecipitated and blotted. (C) RBN2397 inhibits androgen-dependent AR ADP-ribosylation in VCaP prostate cancer cells. AR was immunoprecipitated and protein and ADP-ribose were detected. (D) Dose-response and RBN2397 EC_50_ values for recombinant PARP7 and PARP1 ADP-ribosyltransferase activities measured in vitro, using Histone H2A and H2B as a substrate. (E) Heat maps depicting the effect of RBN2397 (7.6 nM) on the ADP-ribosyltransferase activity of individual PARP family members in vitro (left), and compared to the expression levels of PARP family members in human prostate cancers (TCGA; RNA-seq data) (right). (F) Protein half-life measurements of HA-PARP7 determined in the absence and presence of RBN2397. Cells were pretreated with RBN2397 for 1 hr prior to addition of cycloheximide (0.1 mg/ml). Error bars reflect the standard deviation of biological triplicates. (G) Auto-ADP-ribosylation of ectopically-expressed PARP7 detected in HA-immunoprecipitates from PC3-AR(HA-PARP7) cells treated with RBN2397. Proteasome inhibition with MG132 (10μM, 4 hrs) was used to increase PARP7 levels.

### Stabilization of PARP7 protein

RBN2397 treatment of PC3-AR cells increases the level of PARP7 protein detected by immunoblotting, suggesting the inhibitor might stabilize PARP7 protein (**Fig. 1A**). We performed a time course of cycloheximide treatment and found RBN2397 increases the protein half-life of PARP7 approximately 4-fold (**Fig. 1F**). Stabilization of PARP7 protein provides additional evidence for the on-target effects of RBN2397 in prostate cancer cells. Lastly, we examined the effect of RBN2397 on PARP7 auto-ADP-ribosylation, as automodification is a common biochemical property of PARP enzymes. In this analysis, we determined that MG132 treatment induced the accumulation of ADP-ribosylated PARP7, but automodification as detected by Fl-Af1521 was eliminated by co-treatment with RBN2397 (**Fig. 1G**).

### Inhibition of cell growth

We examined the effects of RBN2397 on growth with cell lines used to model prostate cancer and therapy resistance. VCaP cells encode WT AR and the V7 variant, and are used to model castrate-resistance owing to amplification of *AR*. CWR22Rv1 is an informative model for therapy resistance mediated by *AR* splice variants that lack the ligand binding domain. PC3-AR is a PC3 derivative engineered to express WT AR, which drives a gene expression profile that overlaps other prostate lines[18]. PC3 cells are AR negative, resistant to therapies based on androgen deprivation, highly motile, and used to study prostate cancer metastasis. Overall, treating the four prostate lines with RBN2397 alone had little or no effect on growth (**Fig. 2A**). Reasoning that low PARP7 expression level might limit the effect of RBN2397 on cell growth, we treated cells with androgen (R1881) to activate AR and induce PARP7 expression. We found that RBN2397 was growth inhibitory in androgen-treated VCaP, CWR22Rv1, and PC3-AR cells (**Fig. 2A**). It is well established that androgen treatment is growth-repressive in some prostate cancer lines. R1881 was growth inhibitory in VCaP and PC3-AR cells; RBN2397 further reduced the growth of VCaP and PC3-AR cells, and in CWR22Rv1 cells had greater effects on growth in the presence of R1881 (**Fig. 2A**). Dose response curves for cell growth in the presence of androgen revealed RBN2397 had half-maximal effects in the nanomolar range (**Fig. 2B; Supplementary Table 1**), which is similar to its potency for inhibiting PARP7 enzyme activity in cells (**Fig. 1A**). Growth inhibition by RBN2397 does not involve accumulation of cells in a specific phase of the cell cycle, though within the data a G1 effect of androgen is apparent in VCaP and PC3-AR cells (**Supplemental Fig. 1A**). In VCaP, CWR22Rv1, and PC3-AR cells, RBN2397+R1881 promoted the appearance of rounded cells visible by phase contrast microscopy (**Fig. 2C**), and slightly increased the number of dead cells detected by Trypan blue staining (**Fig. 2D**). Overall, the data suggest that the growth inhibition by RBN2397 includes cytostatic and cytotoxic effects, but these occur without blocking a specific phase of cell cycle to the extent it can be recognized by DNA content.

**Fig.2.**
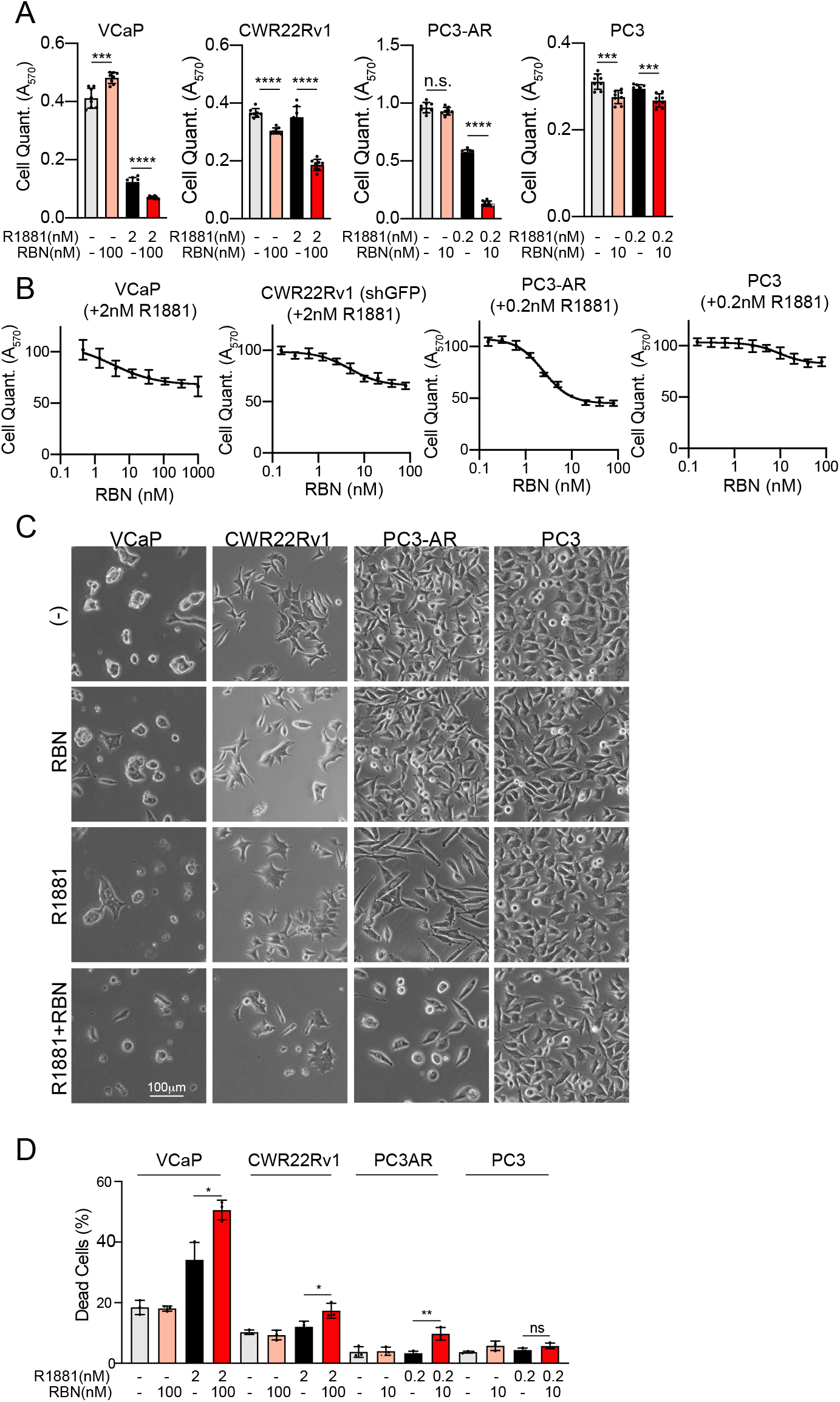
Growth-inhibitory effect of RBN2397 in AR-positive prostate cancer cells in the presence of androgen. (A) Cell growth (MTT assay) in response to combinations of RBN2397 and R1881. Each condition reflects eight biological replicates. For A-C, the error bars depict the standard deviation. ****, p<0.0001; ***, p<0.001; n.s., not significant. (B) RBN2397 dose response growth curves from VCaP, CWR22Rv1, PC3-AR, and PC3 in the presence of the R1881. (C) Phase contrast microscopy showing the morphology of cells in the presence or absence of RBN2397 and R1881 for 6 days (slower growing VCaP and CWR22Rv1) and 3 days (faster growing PC3-AR and PC3). (D) Dead cell detection by Trypan blue staining after culturing cells as in (C). The experiments were performed in triplicate.

We examined the relationship between growth inhibition by RBN2397 and PARP7 expression. Using an siRNA that targets *PARP7* (*siPARP7*), we determined the cell growth reduction caused by RBN2397 in androgen-treated PC3-AR cells was reduced by *PARP7* knockdown (**Fig. 3A**). Endogenous PARP7 protein levels detected by immunoblotting after R1881 treatment are very low, but the additional treatment with RBN2397 stabilizes PARP7 protein and enhances its detection (**Fig. 3A-C**). Reducing PARP7 levels with *shPARP7* (stably-expressed) gave similar results. Though the efficiency of PARP7 depletion and growth differences +RBN2397 in control and *shPARP7* cell lines were quantitively small, the effect of PARP7 knockdown was observed at multiple drug concentrations and was statistically significant in all three cell lines (**Fig. 3A-C**).

**Fig.3.**
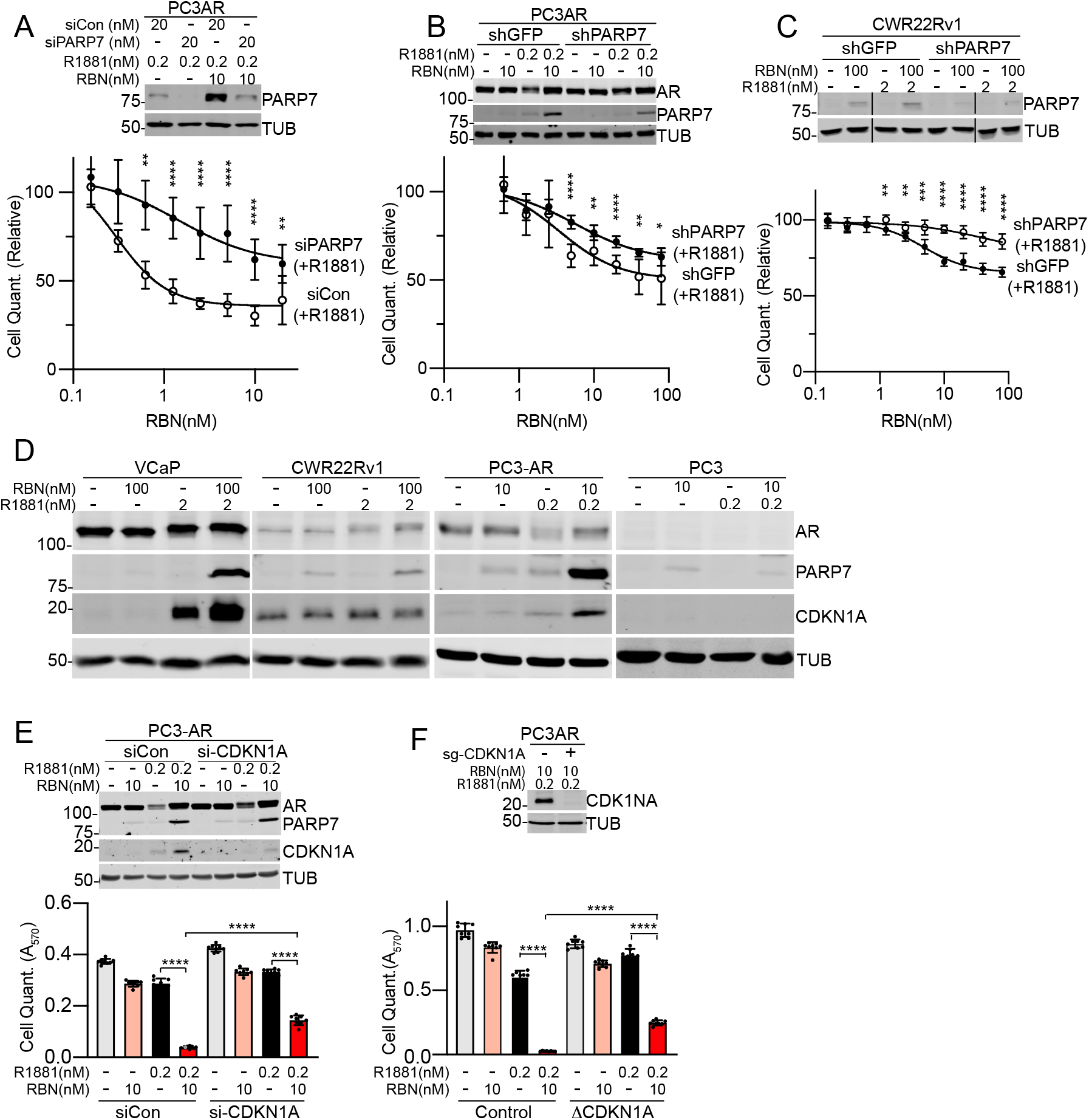
Growth inhibition by RBN2397 is PARP7-dependent and associated with elevated CDKN1A (p21) expression. (A) Treating PC3-AR cells with si*PARP7* partially reduces the growth inhibitory effect of RBN2397. PARP7 knockdown in the cell growth experiments (A-C) was confirmed by immunoblotting for PARP7, with TUBULIN as a loading control (upper panels). For the cell growth in A-C and E-F, each condition reflects eight biological replicates, error bars show the standard deviation, and the statistical differences between cell lines at each RBN concentration are ****, p<0.0001; ***, p<0.001; **, <0.01; *, <0.05 (lower panels). (B, C) Stable knockdown of PARP7 in PC3-AR and CWR22Rv1 cells partially protects from the growth inhibitory effects of RBN2397. CWR22Rv1(*shGFP*) cell growth is also shown in Fig. 2D. (D) Immunoblot detection of CDK inhibitor CDKN1A and PARP7 in prostate cancer lines grown +R1881 and +RBN2397 for 24 hrs. (E) Treating PC3-AR cells with *si-CDKN1A* (20 nM) partially reverses the growth inhibitory effect of RBN2397. The expression levels of CDKN1A and PARP7 +R1881 and +RBN2397 were determined by immunoblotting (upper panel). (F) CRISPR knockout of *CDKN1A* in PC3-AR cells reduces the growth inhibitory effect of RBN2397. The knockout was confirmed by DNA sequencing (not shown) and by immunoblotting for CDKN1A (upper panel).

### CDKN1A (p21) contributes to growth inhibition

One mechanism by which androgens repress the growth of prostate cancer cells through AR is by induction of the CYCLIN-dependent kinase inhibitor CDKN1A (p21) [24]. Given the growth repressive effects of RBN2397 were associated with androgen treatment, we tested whether the RBN2397 effect on growth involves CDKN1A expression. By immunoblotting, CDKN1A protein expression in VCaP and PC3-AR cells is positively regulated by androgen, and CDKN1A levels were increased further by RBN2397 treatment (**Fig. 3D**). The levels of CDKN1A were not affected by androgen in AR-positive CWR22Rv1 cells, and as expected, there was no androgen effect in AR-negative PC3 cells (**Fig. 3D**). Using siRNA to reduce CDKN1A expression, we found that CDKN1A depletion partially rescued the growth inhibitory effect of RBN2397 (**Fig. 3E**). Similarly, deletion of the *CDKN1A* gene in PC3-AR cells (CRISPR-CAS9) was sufficient to achieve a partial rescue of growth in cells treated with RBN2397 (**Fig. 3F**). Thus, in the context of androgen signaling, the growth inhibitory effects of RBN2397 can be partly mediated through CDKN1A expression.

### Exploiting AHR for *PARP7* induction

Growth inhibition by RBN2397 in androgen-treated VCaP, CWR22Rv1, and PC3-AR cells raises the intriguing possibility that PARP7 might be an actionable target in prostate cancer. As an alternative approach for controlling *PARP7* in prostate cancer cells, we turned to the transcription factor AHR, a member of the basic helix-loop-helix /Per-AHR nuclear translocator-Sim protein family. AHR is known to control PARP7 expression in other settings, and has been studied in the context of detoxification mechanisms in liver[7, 25]. AHR is expressed in prostate cancer cells and induces its known target genes *CYP1A1* and *CYP1B1* (**Supplementary Fig. 2A**).

We selected two AHR agonists to explore whether PARP7 induction can sensitize cells to RBN2397 independent of androgen signaling. 6-Formylindolo[3,2-b]carbazole (FICZ) is a naturally occurring tryptophan photoproduct, while 10-Chloro-7*H*-benzimidazo[2,1 -*a*]benz[d*e*]isoquinolin-7-one (10-CI-BBQ; here termed BBQ) is a synthetic molecule. FICZ and BBQ induce AHR target genes through a chaperone-mediated mechanism[26]. FICZ and BBQ (10-100 nM) were both effective for sensitizing cells to growth inhibition by RBN2397 (**Supplementary Table 1**), and the growth inhibitory effect was associated with PARP7 protein stabilization detected by immunoblotting (**Fig. 4A, Supplementary Fig. 2B-C**). These data show that PARP7 can be induced by AHR in AR-positive (CWR22Rv1) and AR-negative (PC3 and DU145) prostate cancer cells for the purpose of sensitizing cells to RBN2397 (**Supplementary Fig. 2B-C**). Combining PARP7 induction with BBQ and RBN2397 treatment reduced the protein levels of CYCLIN A and CYCLIN B in PC3, DU145, and CWR22Rv1, though the effects were small, and not observed in VCaP cells (**Supplementary Fig. 2C**). BBQ and RBN2397 treatment increased the G1 fraction in PC3 and DU145 cells, and slightly reduced the S-phase fraction in CWR22Rv1 cells (**Supplementary Fig. 1B**). Combining BBQ and RBN2397 increased the number of Trypan blue-positive cells in PC3, DU145, and CWR22Rv1 cells (**Fig. 4B**), which is consistent with our finding that there was a reduction in the number of adherent cells in PC3, DU145, and CWR22Rv1 cells treated with BBQ and RBN2397, as detected by phase contrast microscopy (**Fig. 4C**).

**Fig.4.**
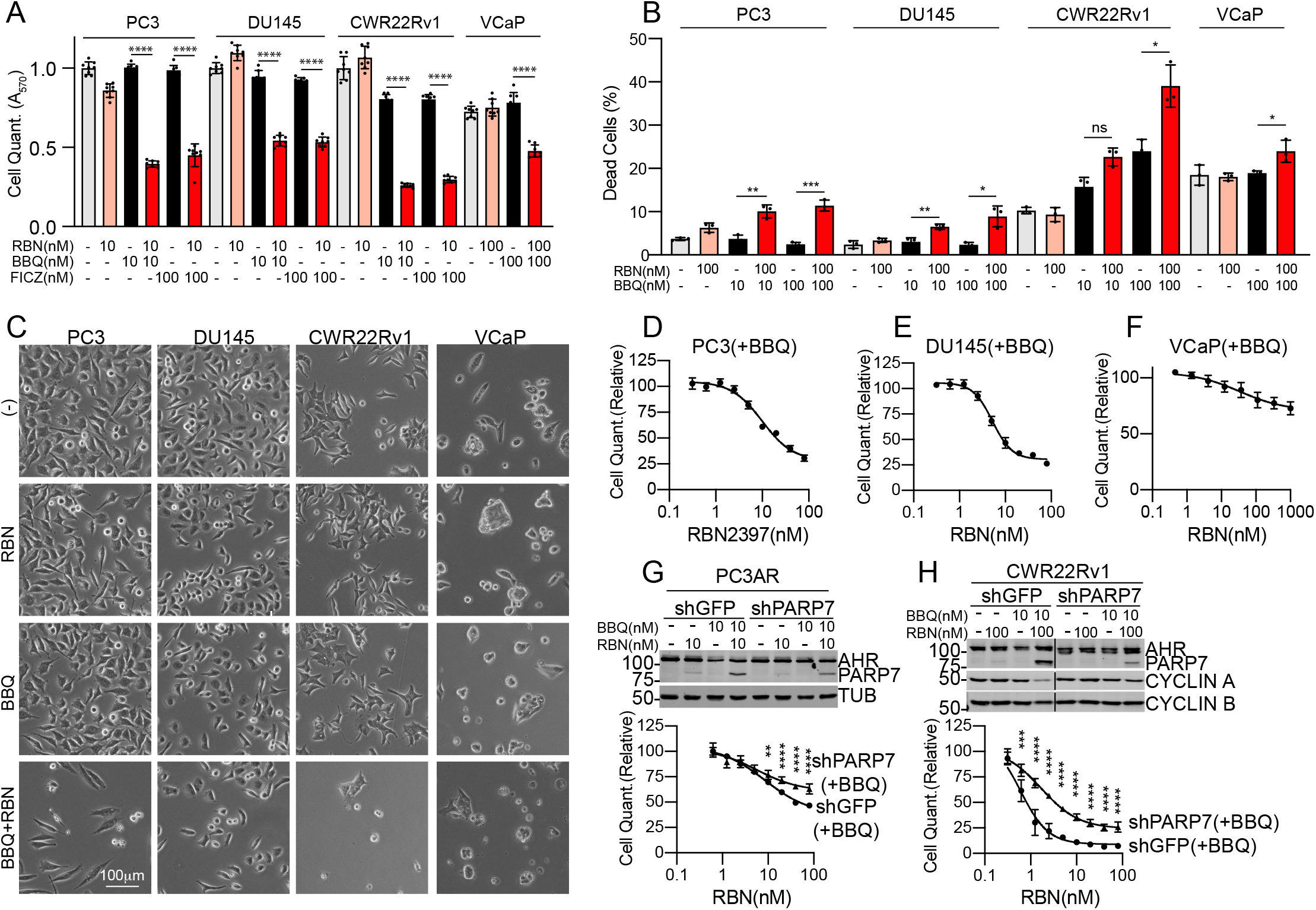
PARP7 induction using aryl hydrocarbon receptor (AHR) agonists sensitizes prostate cancer cells to growth inhibition by RBN2397. (A) Effect of RBN2397 on AR- (PC3, DU145) and AR+ (CWR22Rv1, VCaP) prostate cancer lines treated with AHR agonists BBQ and FICZ to induce PARP7. In panels A, D-H, each condition reflects eight biological replicates, error bars show the standard deviation, and ****, p<0.0001; ***, p<0.001; **, <0.01. (B) Dead cell detection by Trypan blue staining after culturing cells in the presence or absence of RBN2397 and BBQ for 6 days (slower growing VCaP and CWR22Rv1) and 3 days (faster growing PC3-AR and PC3). The experiments were performed in triplicate, error bars show the standard deviation, and ***, p<0.001; **, <0.01, *, <0.05; ns, not significant. (C) Phase contrast microscopy showing the morphology of cells treated with AHR agonist and RBN2397. (D-F) RBN2397 dose response of cell lines grown with BBQ. (G, H) Stable knockdown of PARP7 blunts the growth inhibitory effects of RBN2397 and BBQ co-treatment in PC3-AR and CWR22Rv1 cells.

The cell growth and immunoblotting data for PARP7, CYCLIN A, CYCLIN B, and the Trypan blue detection of dead cells, imply the BBQ and RBN2397 combination is more effective in PC3, DU145, and CWR22Rv1 than in VCaP cells (**Supplementary Table 1**). The RBN2397 dose response curves for cell growth were also consistent with this notion, where PC3 and DU145 cells were inhibited by nanomolar concentrations of RBN2397 and the overall response was greater than VCaP cells (**Fig. 4D-F, H**). To determine if the RBN2397 effect is dependent on PARP7 expression, we compared the RBN2397 sensitivity of control and *PARP7* knockdown cells treated with BBQ. Although the *PARP7* knockdown was partial, the RBN2397 effect was reduced significantly in *shPARP7* compared with *shGFP* cells (**Fig. 4G, H**). Taken together, the data show that RBN2397 can be used to inhibit growth in prostate cancer cells where PARP7 is induced by AHR signaling.

### Growth inhibition is IFN-independent

The cellular effects of RBN2397 in preclinical models of lung cancer were attributed to reversal of a PARP7-TBK1 inhibitory mechanism that restores IFN signaling dependent on IRF3 [12]. Because IRF3 can also regulate the cell cycle [27], we queried whether TBK1 activity and IFN signaling are involved in the growth inhibitory effects of RBN2397 in prostate cancer cells. To this end, we employed specific inhibitors to TBK1 (GSK8612; GSK) and JAK1/2 (Ruxolitnib; Ruxo) in prostate cancer cell lines and induced PARP7 via AHR (PC3, DU145) and AR (PC3-AR)(**Supplementary Fig. 3**). In PC3 cells, RBN2397 treatment increased the basal level of phospho-STAT1, an effect that was eliminated by JAK1/2 (Ruxo) and TBK1 (GSK) inhibitors (**Supplementary Fig. 3B**). These data are consistent with TBK1 acting upstream of JAK1/2 in PC3 prostate cancer cells, as shown in other cell types [12]. Induction of PARP7 by the AHR ligand BBQ does not further increase STAT1 phosphorylation, suggesting the relatively low, basal level of PARP7 expression is sufficient for the RBN2397 effect on STAT1 phosphorylation in PC3 cells (**Supplementary Fig. 3B**, upper panel). This contrasts with the growth inhibitory effect of RBN2397, which is responsive to PARP7 expression induced by treating cells with ligands to AHR and AR (**Figs. 2, 4**).

The growth inhibitory effect of RBN2397 in PC3 cells is not reversed by JAK1/2 and TBK1 kinase inhibition with Ruxo and GSK, respectively (**Supplementary Fig. 3B**, lower panel). This result indicates the RBN2397 enhancement of IFN signaling implied by STAT1 phosphorylation (**Supplementary Fig. 3B**, upper panel) does not drive growth inhibition in PC3 cells. Consistent with this view, induction of IFN signaling only slightly reduces PC3 cell growth (**Supplementary Fig. 3B**, lower panel). In DU145 cells, RBN2397 has no obvious effect on basal STAT1 phosphorylation, despite the fact this line can activate JAK1/2-STAT signaling in response to IFNα (**Supplementary Fig. 3C**, upper panel). Thus, the growth reduction in DU145 cells caused by RBN2397 is not associated with an effect on STAT1 phosphorylation, and blocking JAK1/2 and TBK1 kinase function in this line slightly increases the growth inhibitory effect of RBN2397 (**Supplementary Fig. 3C**, lower panel). In PC3-AR cells, the signaling and growth data are very similar to PC3 cells; RBN2397 increases STAT1 phosphorylation, and blockade of JAK1/2 and TBK1 inhibitors largely eliminates the RBN2397 effect on STAT1 phosphorylation without affecting the growth inhibitory effect of RBN2397 (**Supplementary Fig. 3D**). In VCaP and CWR22Rv1 cells, RBN2397 also has no obvious effect on basal STAT1 phosphorylation (**Supplementary Fig. 3E-F**). From these data, we conclude the growth inhibitory effect of RBN2397 in prostate cancer cells is separable from its effects on IFN signaling that are linked to the kinase activities of JAK1/2 and TBK1.

### RBN2397 Inhibits NCI-H660 Cells

Castrate-resistant prostate cancer can undergo trans-differentiation into neuroendocrine prostate cancer (NEPC), de novo, or in response to potent AR inhibition [28]. We tested the effect of RBN2397 on NCI-H660 cells, which are a widely used cell culture model for NEPC. Because NCI-H660 cells are AR negative, we used the scheme employed with PC3 and DU145 cells (**Fig. 4**) which also lack AR. Co-treatment with AHR agonists and RBN2397 resulted in PARP7 protein accumulation, cell growth inhibition, and fewer cells detected by phase contrast microscopy (**Supplementary Fig. 4A, B**). The RBN2397-mediated growth reduction was associated with a slight increase in apoptotic/necrotic cells, but similar to other cell lines tested, there was not an obvious effect on the cell cycle (**Supplementary Figs. 1C and 5**).

It has been noted that genetic knockout of PARP1 and PARP7 is not the cellular equivalent to drug-induced inhibition of the enzymes[12, 13]. A logical interpretation of these findings is that drug binding to PARP1 and PARP7 exerts dominant negative effects that are in addition to the cellular changes linked to preventing substrate ADP-ribosylation. We considered whether RBN2397 inhibition of prostate cancer cell growth involves a dominant negative effect. An alternative explanation, at least in prostate cancer cells, is that gene expression induced by AR and AHR creates a PARP7 dependency for cell growth. We used the prostate cancer line C4-2b for this analysis because the basal level of PARP7 expression is extremely low in these cells, RBN2397 treatment alone has no effect on growth of these cells, and activation of AHR signaling sensitizes the cells to RBN2397 inhibition (**Supplementary Fig. 4C**).

To test for a dominant negative effect of RBN2397-inhibited PARP7, ectopic PARP7 was stably expressed and detected via an N-terminal Avi-tag (Avi-PARP7; **Supplementary Fig. 4D**). Avi-PARP7 is enzymatically active in these cells since it ADP-ribosylates AR and the modification is blocked by RBN2397 (**Supplementary Fig. 4D**). Expression of Avi-PARP7 was sufficient to confer growth inhibition by RBN2397 (**Supplementary Fig. 4E**). Growth inhibition by RBN2397 in these cells was further enhanced by AHR activation with BBQ, which increases the cellular level of PARP7 by induction of endogenous PARP7. AHR loss was shown to be a mechanism for RBN2397 resistance in H1373 cells, indicating the AHR-PARP7 axis operates in multiple cancer types[29]. The possibility that additional factors regulated through AHR contribute to the RBN2397 vulnerability in prostate cancer cells cannot be excluded.

### PARP7 trapping by RBN2397

Chemical inhibitors to PARP1 exert effects on cells by blocking enzyme function, but also via cytotoxic effects attributed to stabilizing PARP1-chromatin interactions in a process termed trapping[13]. Drug-induced trapping of PARP1 can be detected biochemically by immunoblotting the detergent-resistant chromatin fraction [13]. To test if RBN2397 induces PARP7 trapping in the nucleus, we first examined its subcellular localization by IF microscopy. Treating PC3 cells with AHR agonist and CWR22Rv1 cells with R1881 revealed a nuclear distribution for endogenous PARP7 (**Fig. 5A, B**), consistent with our previous work[23]. Cotreatment with RBN2397 caused a striking increase in the nuclear levels of PARP7 in both cell lines (**Fig. 5A, B**). PARP7 expressed after R1881 treatment is released into the soluble fraction and virtually absent from the insoluble fraction (**Fig. 5C**, lanes 3 and 7). By contrast, the combination of R1881 and RBN2397 results in a pool of PARP7 that partitions to the insoluble fraction (**Fig. 5C**, lanes 4 and 8). In PC3 cells, inducing PARP7 with BBQ to activate AHR also resulted in PARP7 trapping (**Fig. 5D**, lanes 4 and 8). The proportion of PARP7 trapped by RBN2397 treatment is similar to that observed when PARP1 is trapped by Olaparib and Niraparib [13]. To assess if trapping can occur independent of AR and AHR signaling, we examined the effect of RBN2397 on HA-tagged PARP7 distribution. Ectopic HA-PARP7 (stably expressed) underwent RBN2397-induced nuclear trapping detected by microscopy and biochemical fractionation (**Fig. 5E, F**). RBN2397-mediated trapping of PARP7 on chromatin could contribute to the growth inhibition observed in prostate cancer cells.

**Fig. 5.**
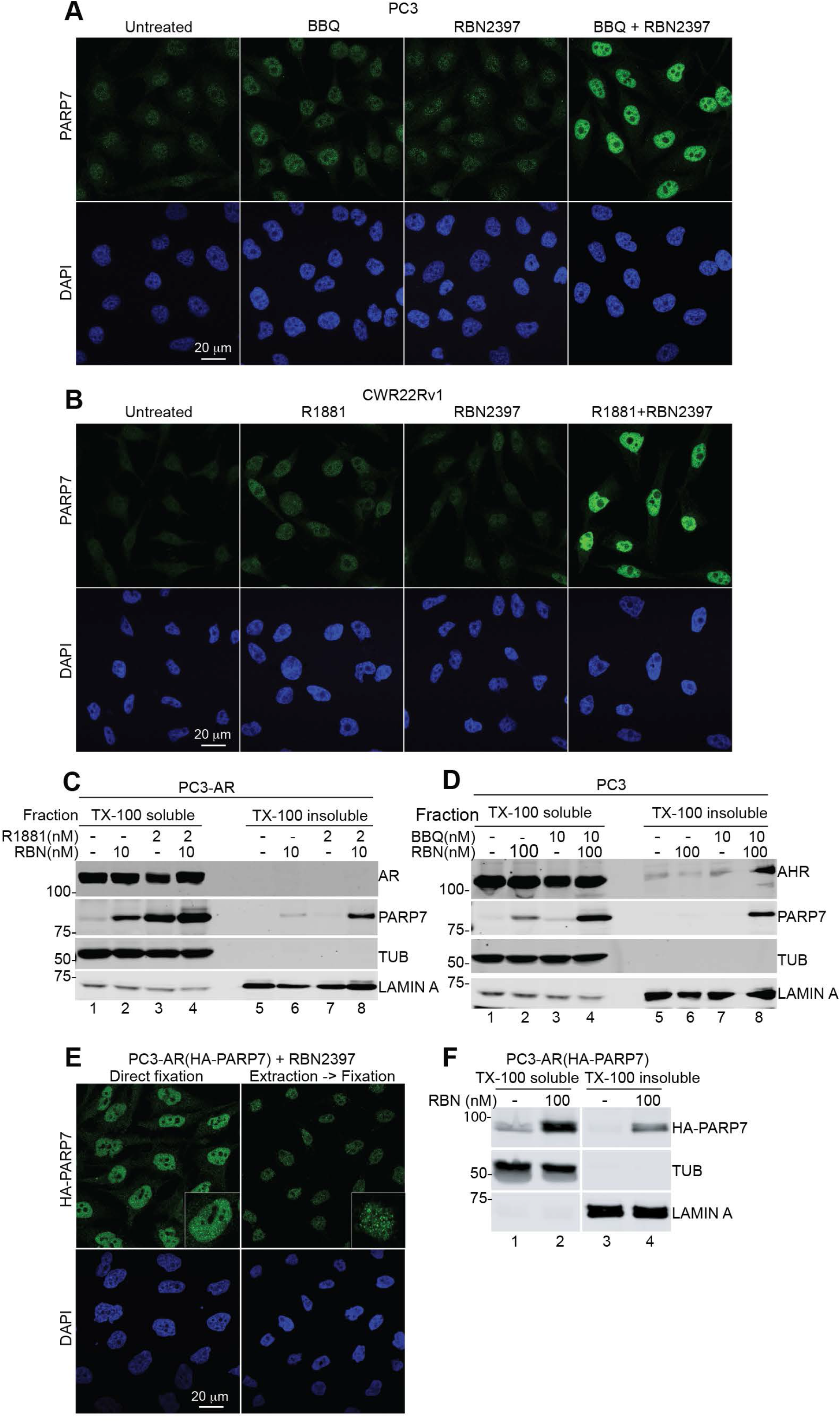
RBN2397 traps PARP7 in the nucleus. (A) IF microscopy of PARP7 in PC3 cells after treatment with RBN2397 and BBQ. (B) IF of PARP7 in CWR22Rv1 after treatment with RBN2397 and R1881. (C) Biochemical fractionation of PC3-AR cells with RBN2397 and R1881 treatment and immunoblot detection of PARP7, AR, TUBULIN and LAMIN A. (D) Biochemical fractionation of PC3 cells treated with RBN2397 and BBQ, and immunoblot detection of PARP7, AHR, TUBULIN and LAMIN A. (E) IF of HA-PARP7 in PC3-AR(HA-PARP7) cells after treatment with RBN2397. The left panel shows cells directly fixed and processed for IF microscopy. The right panel shows cells extracted with TX-100 buffer prior to fixation. (F) Biochemical fractionation of PC3-AR(HA-PARP7) cells with RBN2397 treatment and immunoblot detection of HA-PARP7, TUBULIN and LAMIN A.

### PARP7 Levels in Prostate Cancer

As a first step towards evaluating whether PARP7 levels in human prostate cancer are potentially actionable with RBN2397, we analyzed *PARP7* gene expression data from primary prostate tumors and metastatic *AR*+ and *AR*-prostate tumors[17]. To assess *PARP7* mRNA levels, we used data from the online resource recount3, which uniformly reprocesses publicly available RNA-seq datasets using a Monorail analysis pipeline [30]. This allowed us to compare transcriptomic data from different studies, including cell line and patient samples. We observe there are no changes in *PARP7* mRNA levels in primary tumors (TCGA-PRAD) compared to normal prostate cells (GTEx prostate), but expression is reduced in metastatic tumors [17], and the difference is greater for metastatic tumors that are *AR+* (**Fig. 6A; Supplementary Fig. 6A**). PARP7 expression is slightly higher in ERG fusion positive primary tumors (**Supplementary Fig. 6B**).

**Fig. 6.**
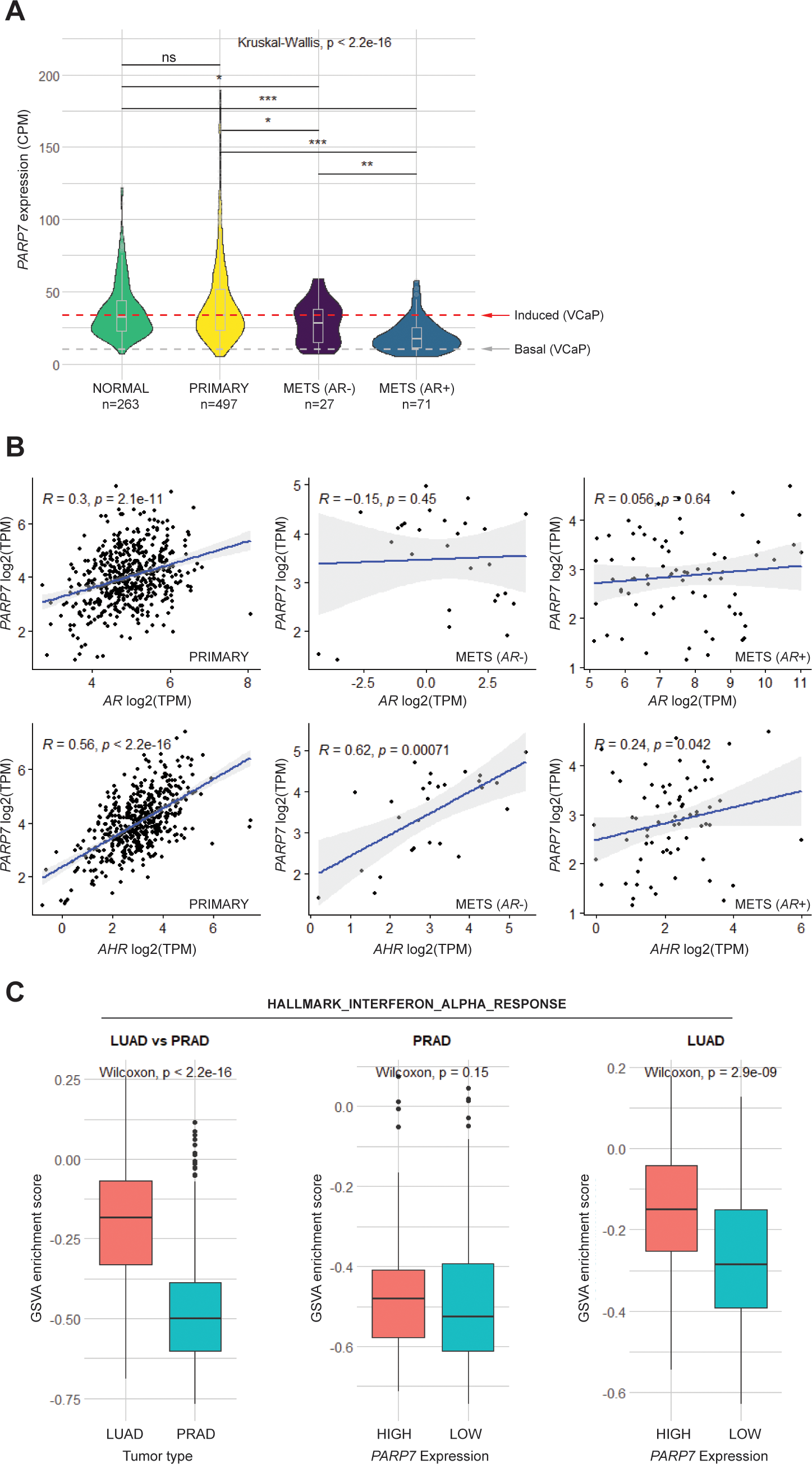
PARP7 expression in prostate cancer. (A) Violin plots (boxplot inserted) showing *PARP7* expression levels in Counts Per Million (CPM) in normal prostate, primary tumors and metastatic AR- and AR+ tumors. The red line indicates the level of *PARP7* in the VCaP cell line required for RBN2397-mediated growth inhibition and the gray line shows the basal level of *PARP7* in VCaP cells. P-values calculated for pairwise comparisons using the Wilcoxon test are indicated (***, <0.001; **, <0.01; *, < 0.05; ns, not significant). All compared values come from the recount3 project. (B) Scatter plots comparing expression of *PARP7* with *AR* and *AHR* in primary tumors (left), metastatic *AR*- tumors (middle), and metastatic *AR*+ tumors (right). Spearman correlation coefficients and p-values are shown on the plots. (C) Boxplots comparing enrichment scores for HALLMARK_INTERFERON_ALPHA_RESPONSE gene set, calculated using GSVA method between lung (LUAD) and prostate (PRAD) primary tumor samples (left); between high *PARP7* expression (top quartile, n=124) and low *PARP7* expression (bottom quartile, n=124) in prostate primary tumor samples (middle); between high *PARP7* expression (top quartile, n=128) and low *PARP7* expression (bottom quartile, n=128) in lung primary tumor samples (right). The p-values from Wilcoxon Rank Sum tests are shown on the plots.

Since growth inhibition by RBN2397 in cell culture is dependent on PARP7 expression, we used our RNA-seq data [18] from VCaP cells treated +R1881 and processed through recount3 to define basal and induced levels of *PARP7* (**Fig. 6A**). We then used the induced level of PARp7 in VCaP cells as a threshold to infer RBN2397 vulnerability in prostate tumors. By this criterion, 50% of primary tumors, 41% of metastatic *AR*-and 11% of *AR*+ tumors are predicted to have PARP7 expression levels that are sufficient for a response to RBN2397. However, this extrapolation will require future validation.

Approximately 1% of primary (4/488) and 8% of metastatic (34/444) prostate tumors show increased genomic copy number in *PARP7*, which could contribute to protein expression levels **(Supplementary Fig. 6C, D, E).** To query how PARP7 expression is potentially regulated in tumors, we computed Spearman’s rank correlation between *PARP7, AR*, and *AHR* gene expression. We found a moderate correlation between *PARP7-AR* expression (r_s_ = 0.30, p < 0.001, n = 497) and a strong positive correlation between *PARP7* and *AHR* expression (r_s_ = 0.56, p < 0.001, n = 497) (**Fig. 6B** left) in primary tumors. In both *AR*+ and *AR*-metastatic tumors the expression of *PARP7* and *AR* is not significantly correlated. However, the *PARP7-AHR* correlation is significant and is especially striking in *AR-* tumors (r_s_ = 0.62, p < 0.001, n = 27) (**Fig. 6B** middle, left). These results suggest that PARP7 expression in primary tumors could be influenced by AR and AHR, but in metastatic tumors the association with AR is lost, whereas the AHR influence is retained. In primary tumors, Gene set variation analysis (GSVA) [20] showed that PARP7 influences the androgen response, but the effect is lost in tumors with low AR expression (**Supplementary Fig. 6F, G, H**).

PARP7 has been proposed to restrain type I IFN signaling in lung cancer models, an effect relieved by RBN2397 [12]. We analyzed the PARP7 influence on type I IFN signaling in primary prostate tumors, using GSVA. We found that lung cancer (TCGA-LUAD) exhibits a much higher enrichment score and pathway activity for genes that are up-regulated in response to IFNα proteins (Type I IFN response) compared to prostate cancer (TCGA-PRAD) (**Fig. 6C**). Moreover, the enrichment scores in LUAD shows an association with *PARP7* expression level, but this is not the case in PRAD (**Fig. 6C**). This suggests that either PARP7 does not influence Type I IFN signaling in primary prostate cancer, or that PARP7 levels in patients are sufficient to saturate the effect.

## DISCUSSION

RBN2397 was developed by optimization of an unselective mono-ADP-ribosyltransferase inhibitor [12]. Biochemical characterization of RBN2397 included using a probe displacement assay to demonstrate potent PARP7 inhibition (IC_50_ <3 nM) and PARP7 selectivity (>50-fold) within the PARP family [12]. These and other data from Ribon Therapeutics [31] demonstrated the potency and selectivity of RBN2397 for PARP7. We characterized the effect of RBN2397 in prostate cancer cells, first by examining its ability to inhibit PARP7 ADP-ribosylation of AR. In prostate cancer cells, AR is a well-defined PARP7 substrate for which the ADP-ribosylation sites have been defined and characterized by mutagenesis[8]. In PC3-AR cells grown without androgen, virtually no ADP-ribose is detected on AR using Fl-Af521 as a probe. Androgen treatment stimulates AR induction of endogenous PARP7, which ADP-ribosylates AR, primarily on Cys sites in the transactivation domain[8]. Biochemical and RNA-seq data are both consistent with PARP7 operating as the androgen-regulated mono-ADP-ribosyltransferase that modifies nuclear AR in prostate cancer cells[8]. In the present study, we found that low nanomolar concentrations of RBN2397 were sufficient to eliminate PARP7 ADP-ribosylation of AR in PC3-AR cells. RBN2397 treatment also prevented ADP-ribosylation of AR in VCaP cells, which demonstrates the effectiveness of RBN2397 in a setting where both PARP7 and its inducer/substrate (AR) are expressed from endogenous genes. Since RBN2397 can also inhibit PARP7 ADP-ribosylation of the estrogen receptor[9], RBN2397 might be useful to target signaling and growth in breast cancer.

A second finding in our study was that RBN2397 can inhibit the growth of prostate cancer cells, and that this occurs under conditions where PARP7 undergoes induced expression. In multiple prostate cancer cell lines, PARP7 can be induced by treating cells with agonists for AR and AHR. The fact that there is little or no growth inhibition by RBN2397 prior to PARP7 induction suggests that the contribution of basal levels of PARP7 to cell growth of these cells in culture is minimal. Consistent with this view, DepMap analysis of VCaP and CWR22Rv1 cells show essentially no dependency on basal PARP7 expression. The growth inhibition by RBN2397 under conditions of PARP7 induction is suggestive of a dominant-negative effect of RBN2397-bound PARP7. Support for a dominant-negative mechanism derives from showing that ectopic PARP7 expression is sufficient to partially sensitize C4-2b cells to the growth inhibition by RBN2397. The growth inhibition of RBN2397 mediated through ectopic PARP7 was not as penetrant as the effect of RBN2397 in androgen-treated cells. The reduced penetrance might be due to an insufficient level of ectopic PARP7, or that androgen signaling through AR affects the expression of genes (in addition to *PARP7*) that promote RBN2397 sensitivity. Basal levels of PARP7 in some cell types such as NCI-H1373 (which display a *PARP7*-dependency DepMap) is sufficient to confer RBN2397 sensitivity [30]. It is tempting to speculate that RBN2397-induced trapping of PARP7 might affect cells through a dominate-negative effect on chromatin-based processes, analogous to the effects of PARP1 inhibitors that block catalytic function-dependent release from chromatin[32]. It is noteworthy that PARP1 trapping can promote IFN signaling through cGAS-STING components, an outcome similar to that reported for RBN2397 inhibition of PARP7 in lung cancer models[12].

In addition to its effect on IFN signaling, RBN2397 treatment of PARP7-expressing cancer cells causes a reduction of cell growth. Whether these effects are linked or separate, and their potential relationship to RBN2397-induced PARP7 trapping reported here remains to be fully explained. Our data favors the conclusion the cell growth and IFN signaling effects in prostate cancer cells are separable effects. RBN2397 enhancement of basal IFN signaling reported by Ribon was shown to occur because PARP7 can restrain TBK1 activity[11, 12]. We found that RBN2397 increases basal IFN signaling in PC3 and PC3-AR cells, as RBN2397 treatment was sufficient to increase phospho-STAT1 levels. The effect on phospho-STAT1 was abrogated by inhibiting either TBK1 or JAK1/2, suggesting the PARP7 effect on TBK1 and IFN signaling in PC3 cells is comparable to some other cell types[12]. And although IFN signaling can be growth repressive, the inhibitory effect of RBN2397 on cell growth in our experiments is not linked to the RBN2397 effect on IFN signaling. Thus, in the aforementioned experiment with TBK1 and JAK1/2 inhibitors that eliminated the RBN2397 enhancement of IFN signaling, blocking TBK1 and JAK1/2 did not alter the RBN2397 inhibition of cell growth. Moreover, RBN2397 had no obvious effect on basal IFN signaling in VCaP, CWR22Rv1, and DU145 cells, despite the fact RBN2397 inhibited growth of these cells. It was noted that RBN2397 can restore basal IFN signaling in CT26 cells without affecting cell proliferation in culture[12]. Whether PARP7 and RBN2397 affect IFN signaling, cell growth, or both, is expected to be dependent on PARP7 expression levels, and also the context. There is an association between PARP7 level and enrichment score for Type 1 IFN signaling in LUAD but not in PRAD. The contribution of PARP7 to IFN signaling might be dependent on the tumor type, but it is also possible that low levels of PARP7 are sufficient for an effect on TBK1. RBN2397 treatment prevents the ADP-ribosylation of >200 proteins based on Af1521 binding[12]. These candidate PARP7 substrates have functions related to nucleic acid metabolism, chromatin structure and transcription. Thus, PARP7 likely participates in a variety of nuclear pathways.

Potential therapeutic benefits of RBN2397 could depend on PARP7 expression levels in tumors, particularly if the growth inhibitory mechanism in tumors involves a dominant negative effect as suggested by our cell culture data. Using recount3, we were able to compare RNA-seq data from cultured cell experiments to data from human prostate tumors. In VCaP cells, growth inhibition by RBN2397 required the addition of agonists for AR or AHR to induce PARP7 expression. Thus, PARP7 levels prior to, and after, androgen treatment of VCaP cells allowed us to define a threshold for PARP7 expression that results in a growth inhibitory response to RBN2397. Using this value as a cutoff, we estimate that about half of all prostate cancers could express sufficient PARP7 to be vulnerable to RBN2397. In *AR*-metastatic tumors, *PARP7* RNA levels are higher on average than *AR*+ tumors; this suggests *AR* status might help predict the utility of RBN2397 in a subgroup of patients. We acknowledge that comparing the expression levels of PARP7 in prostate cancer cells in culture and human prostate tumors is a limitation of the analysis.

In summary, we have shown that PARP7 expression confers sensitivity to growth inhibition by RBN2397 in prostate cancer lines. PARP7 expression levels can be regulated in prostate cancer cells using ligands for AR and AHR, including the NEPC line NCI-H660. PARP7 expression in primary prostate cancer tumors is correlated with both AHR and AR, which is not surprising given the *TIPARP* gene is a direct target of both transcription factors. Interestingly, the positive correlation between *AR* and *PARP7* expression appears to be lost in metastatic prostate tumors, suggesting that *PARP7* becomes highly dependent on AHR signaling for its expression in advanced disease. Based on our cell data, exploiting the AHR pathway and RBN2397 treatment might provide a PARP7-based strategy for inhibiting advanced prostate cancer. While this could include manipulating PARP7 expression with AHR agonists, this approach might be limited by the negative effects of AHR signaling on immune cell function[33]. Levels of the endogenous AHR ligand kynurenine increase with age [34] and with prostate cancer progression[35], which we posit might be sufficient to drive an actionable level of PARP7 expression.

## Author Contributions

**Chunsong Yang** Conceptualization, formal analysis, validation, investigation, visualization, writing–original draft, writing–review and editing; **Krzysztof Wierbiłowicz** Conceptualization, formal analysis, validation, investigation, visualization, writing–original draft, writing–review and editing; **Natalia M Dworak** formal analysis, investigation, visualization; **Song Yi Bae** investigation, formal analysis; **Sachi B. Tengse** investigation, formal analysis; **Nicki Abianeh** investigation; **Justin M. Drake** Conceptualization, formal analysis; **Tarek Abbas** Conceptualization, formal analysis, validation, investigation, writing–original draft; **Aakrosh Ratan** Conceptualization, formal analysis, investigation, visualization, supervision, writing–original draft, writing–review and editing; **David Wotton** Conceptualization, formal analysis, investigation, visualization, writing–original draft, writing–review and editing; **Bryce M Paschal** Conceptualization, supervision, formal analysis, funding acquisition, validation, investigation, visualization, writing–original draft, writing–review and editing.

## ACKNOWLEDGMENTS

We thank Carlos S. Justo-Jaume for technical assistance, Dr. Yuning Jiang for assistance with FACS analysis and Kendall Bromley for comments on the manuscript. We also thank Dr. Heike Keilhack (Ribon Therapeutics) for helpful discussions. The study was supported by NCI award R01CA214872 to B.M.P. and R01GM135376 to T.A.

