## Supplementary Fig. 1-6 for "Induction of PARP7 Creates a Vulnerability for Growth Inhibition by RBN2397 in Prostate Cancer Cells"

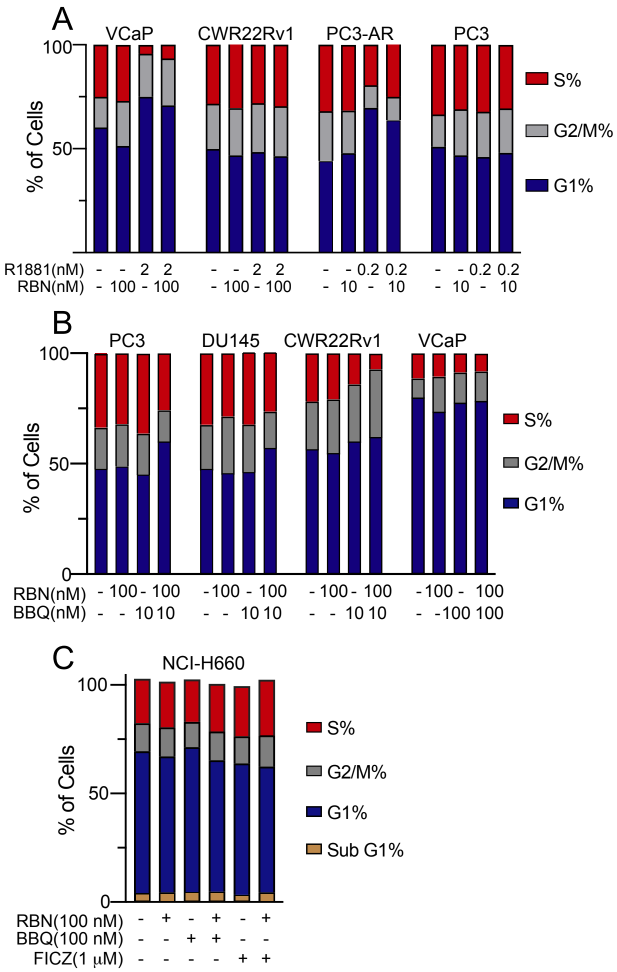


**Supplementary Figure 1. Cell cycle distributions of cell lines treated with RBN2397 plus androgen (A) or plus AHR agonist (B, C).** The treatment durations were 48 hrs for slower growing VCaP, CWR22Rv1 and NCI-H660, and 24 hrs for faster growing PC3-AR and PC3.

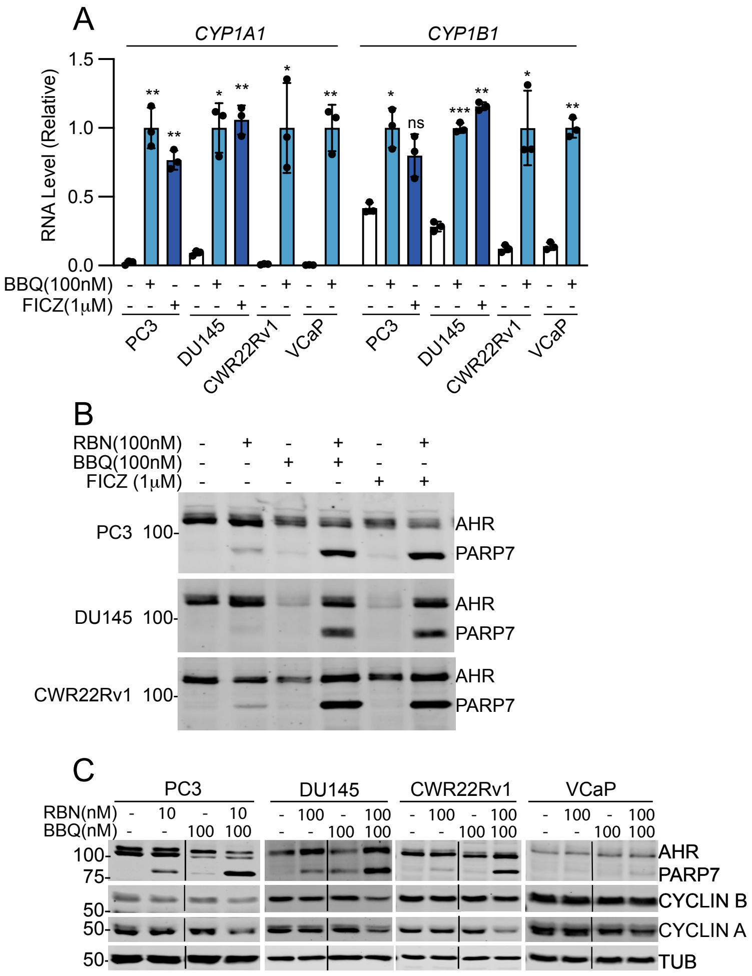


**Supplementary Figure 2. BBQ and FICZ treatment activates AHR signaling in prostate cancer cells.**
(A) *CYP1A1* and *CYP1B1* mRNA levels (normalized to*GUS*) are increased by BBQ and FICZ treatment in the prostate cancer cells. The experiments were done in triplicate. Error bars represent standard deviation. Statistical difference is between the vehicle control and AHR agonist-treated samples. ***, <0.001; **, <0.01; *, <0.05; ns, not significant.
(B) PARP7 protein levels are increased by the treatments with RBN2397 and AHR agonists.
(C) RBN2397 and BBQ combination treatments induce PARP7 protein levels but decrease CYCLIN A protein levels.

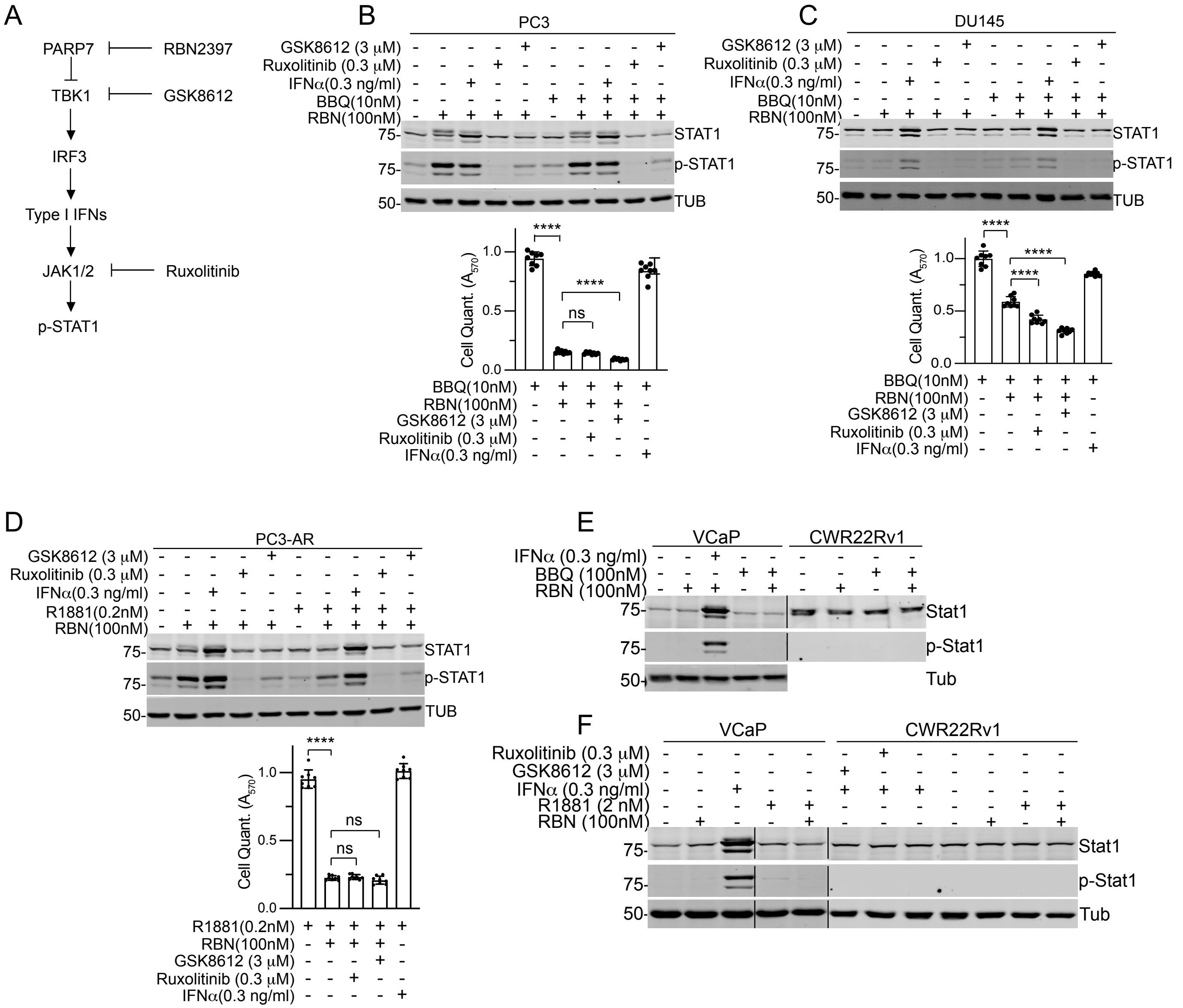


**Supplementary Figure 3. The growth inhibitory effect of RBN2397 in prostate cells is not dependent on TBK1 and JAK1/2 kinase activity.**

(A) The diagram shows a potential mechanism to link RBN2397 to JAK/STAT signaling in cell types including PC3 and PC3-AR cells. pathway of RBN2397 effects to STAT1 signaling in some cells, including PC3 and PC3-AR cells.

(B) Immunoblot detection of STAT1 and Phospho-STAT1 (left) and cell growth assays (right) in PC3 cells treated as indicated. Ruxolitinib (Ruxo, 0.3 μM) and GSK8612 (GSK, 3 μM) were used to inhibit TBK1 and JAK1/2, respectively, and IFNα (0.3 ng/ml) was used to activate JAK/STAT signaling. PARP7 induction and inhibition was mediated by addition of BBQ and RBN2397, respectively.

(C) Phospho-STAT1 levels are not altered detectably by RBN2397 treatment of DU145 cells, and the growth inhibitory effects of RBN2397 are not rescued by blocking TBK1 (GSK) and JAK1/2 (Ruxo) activity. The treatments in DU145 and the analysis were the same as described for panel A.

(D) The same treatments and analysis were as described for panel A, except R1881 was used to induce PARP7 in PC3-AR cells.

(E, F) RBN2397 does not induce STAT1 phosphorylation in VCaP and CWR22Rv1 cells.

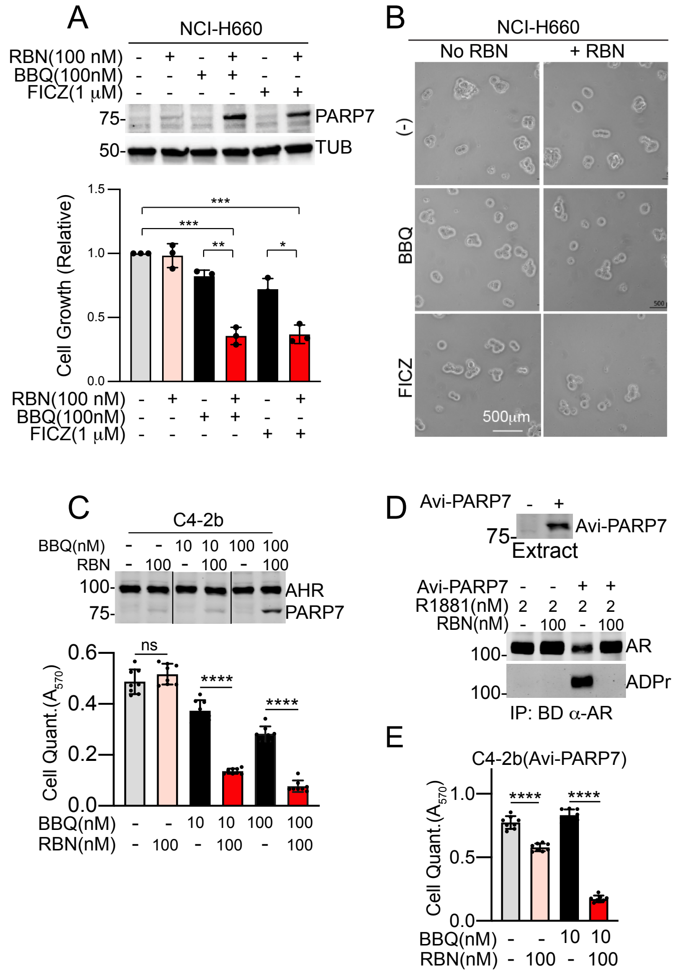


**Supplementary Fig 4. RBN2397 inhibition of prostate cancer cell growth requires PARP7 over-expression.** (A) Effect of RBN2397 on NCI-H660 cells treated with AHR agonists BBQ or FICZ to induce PARP7. PARP7 levels are shown (upper panel). Each condition reflects five biological replicates. In A, C and E, error bars show the standard deviation, and ****, $p<0.0001$; ***, $p<0.001$; **, $<0.01$, *, $p<0.05$; ns, not significant.

(B) Phase contrast microscopy showing the morphology of cells using the treatments shown in (A).

(C) Effect of RBN2397 on C4-2b cell lines treated with AHR agonists BBQ. In C and E, each condition reflects eight biological replicates.

(D) Avi-tagged PARP7 (Avi-PARP7) over-expression in C4-2b cells (upper panel) characterized by immunoprecipitation, immunoblotting, and detection of AR ADP-ribosylation with Fl-Af1521.

(E) RBN2397 inhibited the cell growth of the C4-2b cells over-expressing Avi-Parp7 in the absence of AHR agonist.

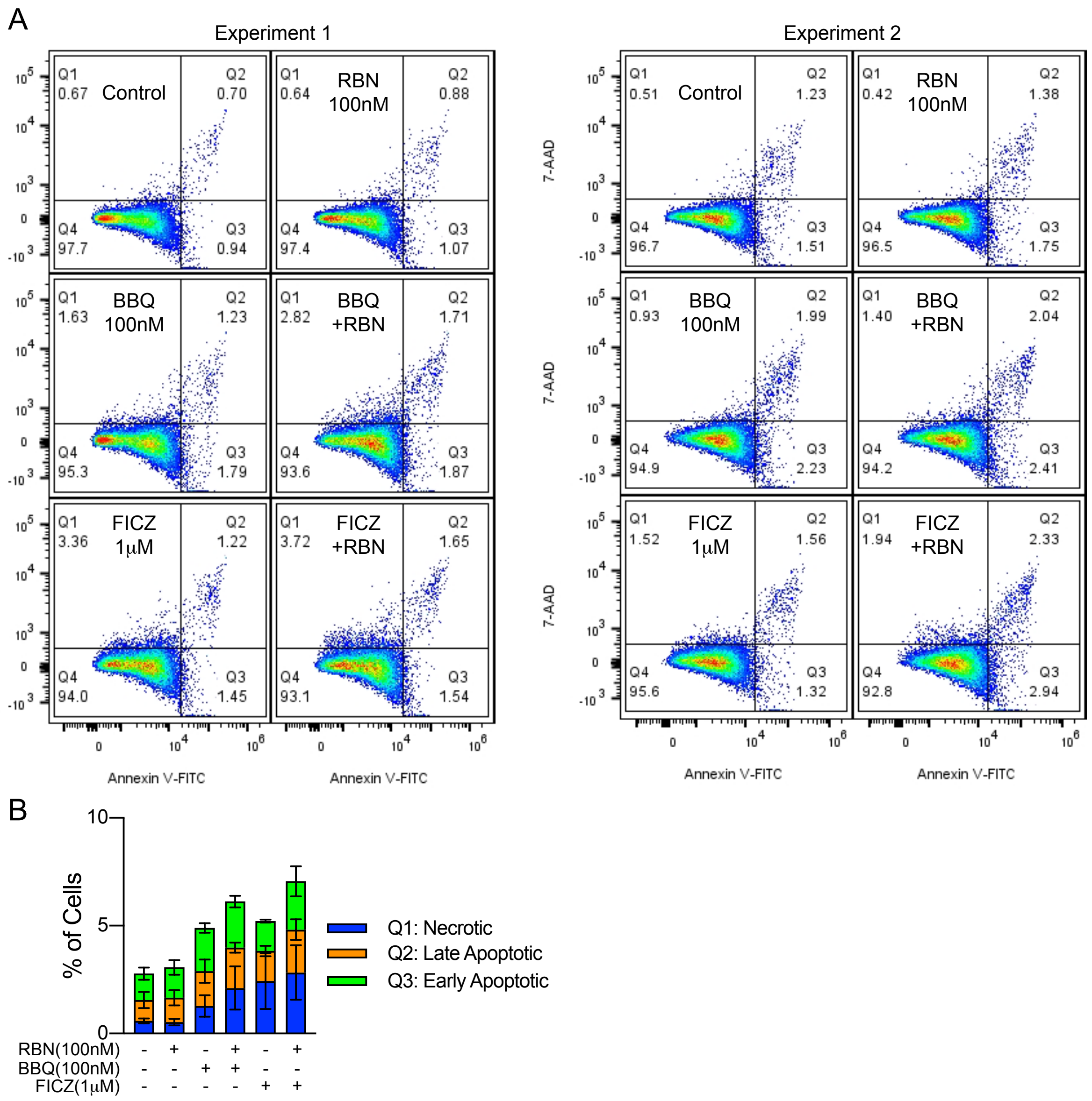


**Supplementary Figure 5. FACS analysis of NCI-H660 cells for detection of necrosis and apoptosis.**
(A) Cells were treated with BBQ, FICZ and RBN for 5 days, stained with Annexin V-FITC and 7-AAD, and examined by flow cytometry.
(B) Percentage of cells in necrosis, late apoptosis and early apoptosis from two experiments were combined and plotted as the mean and range.

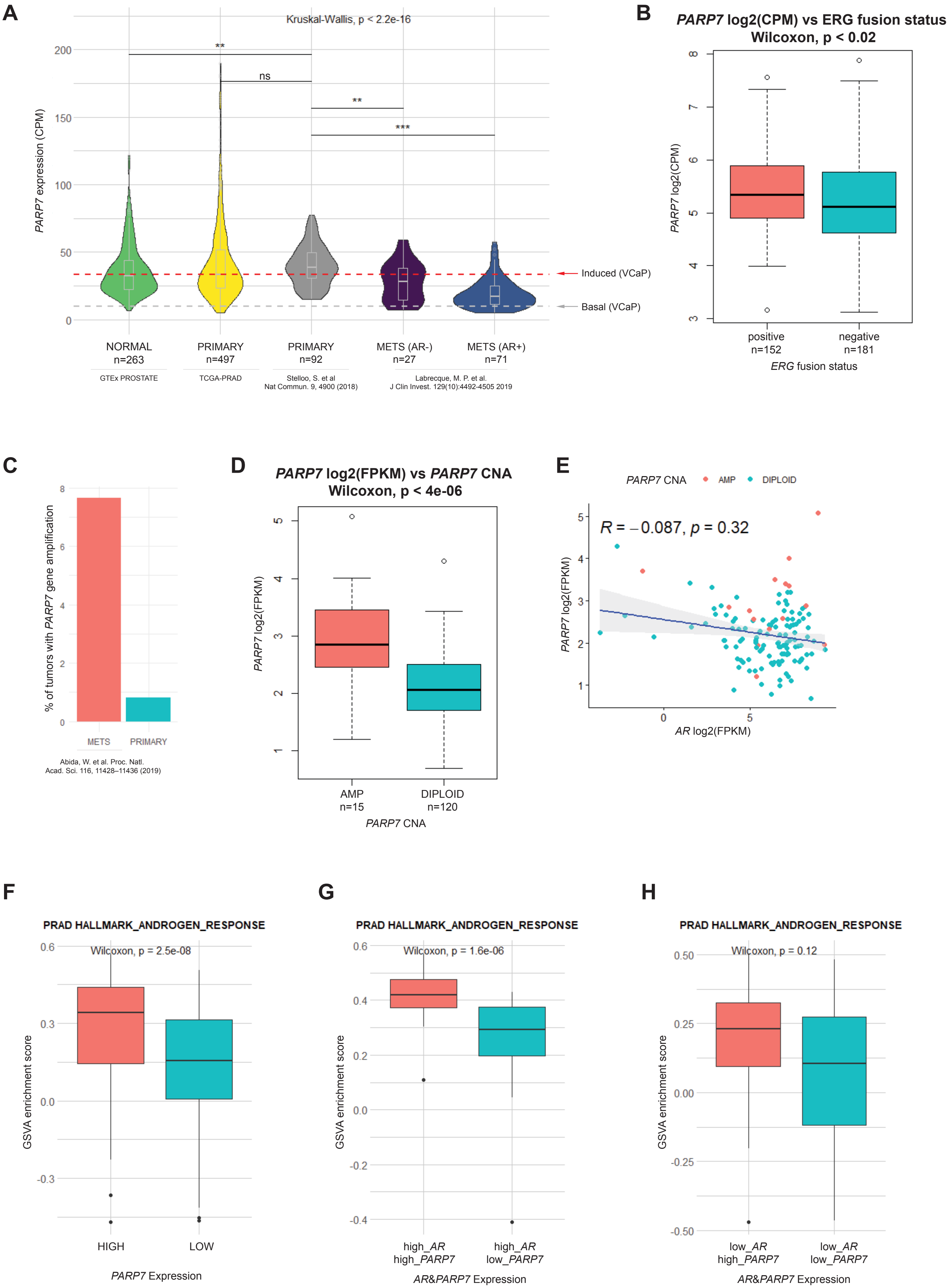
A
Kruskal-Wallis, p < 2.2e-16
PARP7 expression (CPM)
**
ns
**
***
Induced (VCaP)
Basal (VCaP)
NORMAL n=263
PRIMARY n=497
PRIMARY n=92
METS (AR-) n=27
METS (AR+) n=71
GTEx PROSTATE
TCGA-PRAD
Stelloo, S. et al Nat Commun. 9, 4900 (2018)
Labrecque, M. P. et al. J Clin Invest. 129(10):4492-4505 2019
B
PARP7 log2(CPM) vs ERG fusion status
Wilcoxon, p < 0.02
PARP7 log2(CPM)
positive n=152
negative n=181
ERG fusion status
C
% of tumors with PARP7 gene amplification
METS
PRIMARY
Abida, W. et al. Proc. Natl. Acad. Sci. 116, 11428–11436 (2019)
D
PARP7 log2(FPKM) vs PARP7 CNA
Wilcoxon, p < 4e-06
PARP7 log2(FPKM)
AMP n=15
DIPLOID n=120
PARP7 CNA
E
PARP7 CNA
AMP
DIPLOID
R = –0.087, p = 0.32
PARP7 log2(FPKM)
AR log2(FPKM)
F
PRAD HALLMARK_ANDROGEN_RESPONSE
Wilcoxon, p = 2.5e-08
GSVA enrichment score
HIGH
LOW
PARP7 Expression
G
PRAD HALLMARK_ANDROGEN_RESPONSE
Wilcoxon, p = 1.6e-06
GSVA enrichment score
high_AR high_PARP7
high_AR low_PARP7
AR&PARP7 Expression
H
PRAD HALLMARK_ANDROGEN_RESPONSE
Wilcoxon, p = 0.12
GSVA enrichment score
low_AR high_PARP7
low_AR low_PARP7
AR&PARP7 Expression

**Supplementary Figure 6. Extendend analysis of *PARP7* expression in prostate cancer**

(A) Violin plots (boxplots inserted) with the *PARP7* expression data in Counts Per Million (CPM) from Figure 8A, with the additional *PARP7* expression data (grey shading), from Stelloo, S. et al. Nat Commun. 2018. *PARP7* expression is similar in the two primary prostate cancer tumor data sets. The red line indicates the *PARP7* level in VCaP cells induced by R1881 and found to sensitize cells to RBN2397-mediated growth inhibition, and the gray line shows the basal level of PARP7 in VCaP cells. P-values calculated for pairwise comparisons using the Wilcoxon Rank Sum test are indicated (***, <0.001; **, <0.01; *, < 0.05; ns, not significant). These comparisons were done only for additional dataset, since the rest is shown on the fig. 8A. All compared values come from the recount3 project.
(B) Boxplots comparing *PARP7* expression levels (CPM, log2 transformed) in *ERG* fusion positive and negative primary prostate cancers (TCGA-PRAD). P-value calculated using Wilcoxon Rank Sum test is indicated on the figure.
(C) Bar plot showing the percentage of tumors with *PARP7* gene amplification in primary prostate cancer tumors (TCGA-PRAD) and metastatic prostate cancer tumors (SU2C/PCF Dream Team, PNAS 2019). Data was acquired from cBioPortal (https://www.cbioportal.org/).
(D) Boxplots comparing *PARP7* expression level Fragments Per Kilobase Million, FPKM (log2 transformed) in metastatic prostate cancer tumors (SU2C/PCF Dream Team, PNAS 2019) with (AMP) and without (DIPLOID) *PARP7* gene amplification. Data was acquired from cBioPortal (https://www.cbioportal.org/). The data shown reflects the subset of tumors for which RNA-seq data is available. P-value calculated using Wilcoxon Rank Sum test is indicated on the figure.
(E) Scatter plot comparing expression of *PARP7* with AR (FPKM, log2 transformed) in metastatic prostate cancer tumors (SU2C/PCF Dream Team, PNAS 2019). Spearman correlation coefficient and the p-value are shown on the plot. Tumors with (AMP) and without (DIPLOID) *PARP7* gene amplification are labeled as indicated in the legend. Data was acquired from cBioPortal (https://www.cbioportal.org/).
(F) Boxplots comparing enrichment scores for the HALLMARK_ANDROGEN_RESPONSE gene set, calculated using the GSVA method between high *PARP7* expression (top quartile, n=124) and low *PARP7* expression (bottom quartile, n=124) in primary prostate cancer tumors (TCGA-PRAD). P-value calculated using Wilcoxon Rank Sum test is indicated on the figure.
(G) Boxplots comparing enrichment scores for the HALLMARK_ANDROGEN_RESPONSE gene set, calculated using the GSVA method between high *AR* expression (top quartile), high *PARP7* expression (top quartile) (n=32) and with high *AR* expression (top quartile), low *PARP7* expression (bottom quartile) (n=26) in primary prostate cancer tumors (TCGA-PRAD). P-value calculated using Wilcoxon Rank Sum test is indicated on the figure.
(H) Boxplots comparing enrichment scores for the HALLMARK_ANDROGEN_RESPONSE gene set, calculated using the GSVA method between low *AR* expression (bottom quartile), high *PARP7* expression (top quartile) (n=26) and with low *AR* expression (bottom quartile), low *PARP7* expression (bottom quartile) (n=46) in primary prostate cancer tumors (TCGA-PRAD). P-value calculated using Wilcoxon Rank Sum test is indicated on the figure.
