## Supplementary table 1 for "Induction of PARP7 Creates a Vulnerability for Growth Inhibition by RBN2397 in Prostate Cancer Cells"

**Supplementary Table 1. RBN2397 growth effects in prostate cencer cell lines.**

| Cell Line | AR Status | Ligand* (Conc.) | EC50 of RBN2397 (nM) | Growth Inhibition at RBN2397 (nM) |
|---|---|---|---|---|
| VCaP | WT and V7 | R1881 (2 nM) | 3.4 | 28.7% (111 nM) |
| PC3 | (-) | R1881 (0.2 nM) | 9.4 | 15.6% (80 nM) |
| PC3AR | WT | R1881 (0.2 nM) | 2.4 | 54.9% (80 nM) |
| VCaP | WT and V7 | BBQ (100 nM) | 34.2 | 19.8% (111 nM) |
| PC3 | (-) | BBQ (10 nM) | 10 | 69.4% (80 nM) |
| DU145 | (-) | BBQ (10 nM) | 5.2 | 73.6% (80 nM) |
| PC3AR(shGFP) | WT | R1881 (0.2 nM) | 3.4 | 49.0% (80 nM) |
| PC3AR(shParp7) | WT | R1881 (0.2 nM) | 5.7 | 37.0% (80 nM) |
| PC3AR(shGFP) | WT | BBQ (10 nM) | 8.4 | 62.0% (80 nM) |
| PC3AR(shParp7) | WT | BBQ (10 nM) | 5.6 | 39.2% (80 nM) |
| CWR22Rv1(shGFP) | H874Y and V7 | R1881 (2 nM) | 4.8 | 34.4% (80 nM) |
| CWR22Rv1(shPARP7) | H874Y and V7 | R1881 (2 nM) | 39.6 | 14.2% (80 nM) |
| CWR22Rv1(shGFP) | H874Y and V7 | BBQ (10 nM) | 0.6 | 91% (80 nM) |
| CWR22Rv1(shPARP7) | H874Y and V7 | BBQ (10 nM) | 1.7 | 75.8% (80nM) |

*R1881: synthetic androgen; BBQ: AHR agonist 10-Cl-BBQ.
